# Gonadal sex patterns p21-induced cellular senescence in mouse and human glioblastoma

**DOI:** 10.1101/2021.06.02.446756

**Authors:** Lauren Broestl, Lucia Grandison, Saraswati Shenoy, Miranda M. Tallman, Gina Rhee, Wei Yang, Jasmin Sponagel, Najla Kfoury-Beaumont, Cameron M. Hill, Diane D. Mao, Albert H. Kim, Sheila A. Stewart, Monica Venere, Jingqin Luo, Joshua B. Rubin

**Author notes:** Author for correspondence Joshua B. Rubin, MD, PhD CB 8208, 660 South Euclid Ave. St. Louis, MO 63110.

## Abstract

Males exhibit higher incidence and worse prognosis for the majority of cancers, including glioblastoma (GBM). Disparate survival may be related to sex-biased responses to treatment, including radiation. Using a mouse model of GBM, we show that female cells are more sensitive to radiation, and that senescence represents a major component of the radiation therapeutic response in both sexes. Correlation analyses revealed that the CDK inhibitor p21 and irradiation induced senescence were differentially regulated between male and female cells. Indeed, female cellular senescence was more sensitive to changes in p21 levels, a finding that was observed in both wildtype and transformed murine astrocytes and patient-derived GBM cell lines. Using a novel Four Core Genotypes model of GBM, we further show that sex differences in p21-induced senescence are patterned by gonadal sex. These data suggest that sex differences in p21 induced senescence contribute to the female survival advantage in GBM.

## Introduction

Sex differences are observed in the majority of diseases, including cancer. Across a wide range of ages and cultures, male sex is associated with higher incidence and worse prognosis for most tumor types^1–3^. Glioblastoma (GBM), the most common primary malignant brain tumor, follows this same pattern – women are both less likely to develop GBM and have a significant survival advantage compared to men^4–6^. Sex differences in survival may result from differences in the response to standard of care therapy, which for GBM includes surgical resection, followed by treatment with radiation and chemotherapy. Previous research from our lab found that female GBM patients had a greater decline in tumor growth velocity after treatment with radiation and chemotherapy than male patients, and that when these measures were used to stratify patients, there was a significant association with survival in female but not male patients^7^. Furthermore, we identified unique molecular pathways associated with improved survival in male and female patients, and expression of genes in these pathways correlated with the sensitivity to a range of chemotherapeutic agents^7^. While this study advanced our understanding of the relationship between chemotherapy and cellular responses in male and female GBM, it did not focus on radiation, the backbone of GBM treatment, which is currently applied to male and female patients equally.

In cancer treatment, the goal is to stop tumor cell proliferation. While one mechanism to achieve this is through triggering cell death/apoptosis, another possibility is to activate cellular senescence – a cell fate decision that results in irreversible cell cycle arrest^8^. Senescence is a known outcome of radiation treatment, and at least one study has reported that this is the dominant response in glioblastoma^9^. Senescence is often described as a double-edged sword, since senescent cells secrete a broad variety of factors with complex pro- and anti-tumorigenic effects^10–12^. However, the cell intrinsic aspect of senescence, specifically the terminal cell cycle exit of cells harboring mutated DNA, serves a beneficial function in tumor prevention and treatment, and enhancing this response could be a strategy to improve treatment efficacy.

Senescence is primarily regulated by two central pathways: p53/p21^WAF1/Cip1^ and p16^INK4A^/Rb^8, 10, 11^. Importantly, we have previously identified sex differences in the regulation of p21, p16, and RB in a GBM model, with female cells being more likely to upregulate these pathways in response to cellular stress^13, 14^. Whether these differences influence the cellular response to radiation in males and females is currently unknown. In this study, we show that primary male and female human GBM lines have unique molecular pathways that contribute to radiation sensitivity. Using an *in vitro* mouse model of GBM, we find that females are more sensitive to radiation, and that senescence is a major component of the radiation response in both males and females. Using a correlation-based approach, we identify an association between p21 and irradiation induced cellular senescence that differs in males and females. The same levels of p21 correlate with higher percentages of senescent cells in females than in males – a finding that was observed in both wildtype and transformed cells, as well as in patient-derived GBM cell lines. Finally, using a novel Four Core Genotypes (FCG) model of GBM, we investigate the biological mechanisms underlying this sex difference, and show that sex differences in p21-induced senescence are patterned by gonadal sex.

## Results

### Unique molecular pathways contribute to radiation response in male and female primary human GBM cell lines

Building upon our previously published work, which identified unique gene signatures associated with sensitivity to a range of chemotherapeutic agents in male and female primary human GBM cell lines^7^, we sought to determine whether these pathways similarly influenced sensitivity to radiation. We performed irradiation dose response curves with four male and five female primary human GBM lines (Supplementary Table 1; Supplementary Fig. 1), and calculated an individual IC_50_ value for each line. As previously observed for other DNA damaging agents, there was no significant difference in the median IC_50_ values for male and female lines (Fig. 1a).

**Fig. 1:**
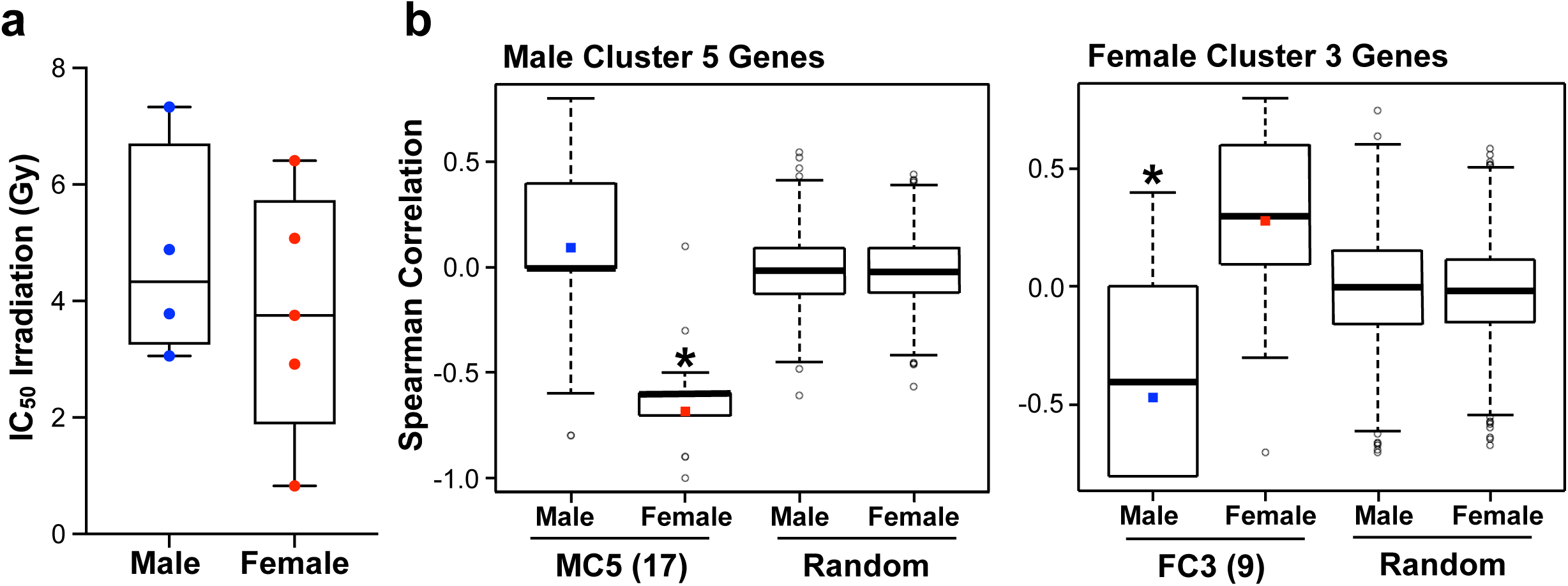
Unique pathways contribute to cell-intrinsic radiation sensitivity in male and female human GBM lines. **a** Box plots of irradiation IC_50_ values for four male and five female primary human GBM lines (horizontal bar indicates median). **b** Spearman correlation coefficients of irradiation IC_50_ values with expression of MC5 genes (17 genes), FC3 genes (9 genes), or random gene sets of the same size. For MC5 and FC3, box plots represent the distribution of the correlation coefficients for the 17 and 9 genes respectively. The colored square marks the average. For the random gene sets, the box plots represent the distribution of the Olkin-averaged Spearman correlation coefficient for 1000 gene sets of 17 (Male Group5) or 9 (Female Group3) randomly selected genes. MC5: Male p=0.245, Female p<0.001. FC3: Male p=0.025, Female p=0.083.

**Fig. 2:**
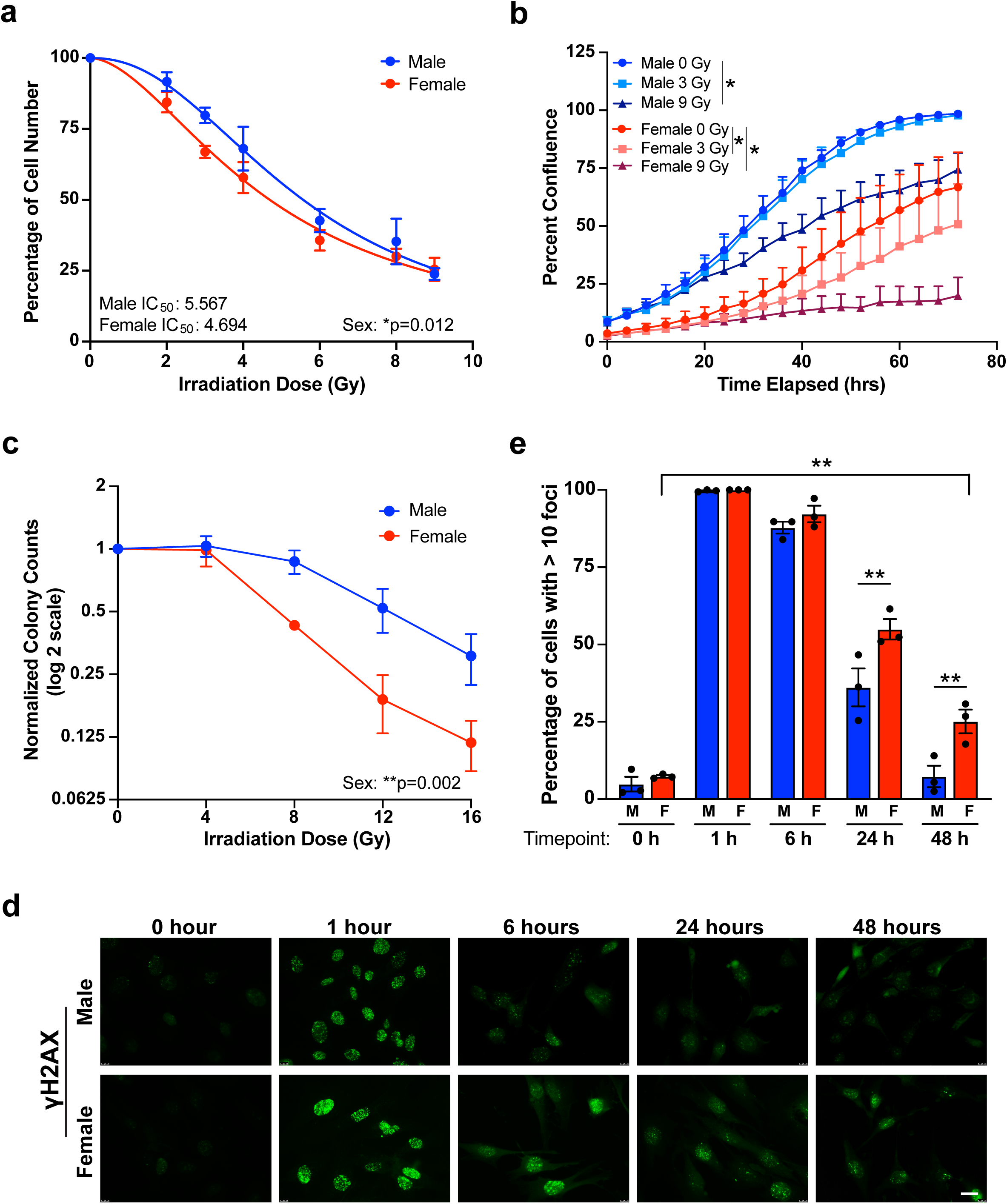
Female mouse GBM model astrocytes are more sensitive to radiation treatment. **a** Irradiation dose response curves in male and female *Nf1-/- DNp53* astrocytes. Curves represent combined values from five separate established cell lines – each consisting of a corresponding male and female line generated at the same time. Male IC_50_ = 5.567 (95% CI 5.094-6.088), Female IC_50_ = 4.694 (95% CI 4.349-5.061). Two-way ANOVA: Dose p<0.0001, Sex p=0.0123, Interaction p=0.6034. Data are means +/-SEM (n=5/sex). **b** Male and female *Nf1-/- DNp53* cell growth curves after irradiation with 0, 3, or 9 Gy. Cell growth was tracked using live cell imaging; images were taken every 4 hours for 3 days and used to calculate percent confluence over time. Curves represent combined values from three male and three female cell lines. Two-way repeated measures ANOVA – Male 0 Gy vs Male 3 Gy: Time p<0.0001, Dose p=0.3394, Interaction p=0.9087; Female 0 Gy vs Female 3 Gy: Time p<0.0001, Dose p=0.0183, Interaction p<0.0001; Male 0 Gy vs Male 9 Gy: Time p<0.0001, Dose p=0.0325, Interaction p<0.0001; Female 0 Gy vs Female 9 Gy: Time p<0.0001, Dose p=0.05, Interaction p<0.0001. Data are means +/-SEM (n=3/sex). **c** Normalized colony counts from the clonogenic assay in male and female *Nf1-/- DNp53* cells 5 days after irradiation with 0, 4, 8, 12, or 16 Gy. Two-way ANOVA: Dose p<0.0001, Sex p=0.002, Interaction p=0.111. Data are means +/-SEM (n=3/sex). **d** Representative immunofluorescence images of male and female *Nf1-/- DNp53* astrocytes stained for γH2AX 0, 1, 6, 24, and 48 hours after irradiation with 3 Gy. Scale bar, 20 μm. **e** Quantification of the percent of cells with >10 γH2AX foci at each timepoint in male and female *Nf1-/- DNp53* astrocytes following irradiation with 3 Gy. Two-way ANOVA: Time p<0.0001, Sex p=0.0002, Interaction p=0.0121. **p<0.01 as indicated by bracket. Data are means +/-SEM (n=3/sex).

Our previous work identified a set of differentially regulated genes associated with better survival in either male (Male Cluster 5 – MC5) or female (Female Cluster 3 – FC3) GBM patients (see Fig. 3A, B and Fig. 5C, D in reference^7^). Pathway analysis of these gene sets identified enrichment of a cell cycle regulation pathway in males (MC5 – 17 genes) and an integrin signaling pathway in females (FC3 – 9 genes) (Supplementary Table 2). In both cases, downregulation of the majority of genes in the pathway was associated with improved survival^7^. We previously observed that low expression of the MC5 gene set was associated with low IC_50_ for multiple chemotherapies in male cell lines, while low expression of the FC3 gene set was associated with low IC_50_ for multiple chemotherapies in female cell lines – a finding consistent with the observation that downregulation of the MC5 and FC3 genes was associated with improved survival in male or female GBM patients respectively^7^. To determine whether these molecular pathways may also be contributing to the male and female irradiation response, we calculated Spearman rank correlation coefficients between IC_50_ values and the expression levels for MC5 and FC3 genes, measured using the Illumina HumanHT-12 v4 expression microarray. For the female cell lines, we saw a similar pattern to that observed with other DNA damaging agents (see Fig. 7B, C in reference^7^). There was a mild positive correlation between FC3 gene expression and irradiation IC_50_, indicating that low expression of these genes was associated with low IC_50_, or better response to irradiation. There was also a negative correlation between MC5 gene expression and IC_50_, indicating that high expression of the MC5 genes was associated with low IC_50_ (i.e. better response to irradiation) in the female lines (Fig. 1b). In contrast, the male cell lines showed no significant correlation between MC5 gene expression and IC_50_ in the male lines, while there was a significant negative correlation between FC3 gene expression and IC_50_, indicating that high expression of the FC3 genes was associated with better response to irradiation in the male lines (Fig 1b). Neither male nor female lines showed any significant correlation between IC_50_ and the expression levels of randomly selected gene sets, serving as a negative control. These results suggest that the response to radiation therapy may be driven by different intracellular pathways in males and females, and that efforts to improve response to therapy may be advanced by identifying the mechanism(s) underlying this sex difference.

**Fig. 3:**
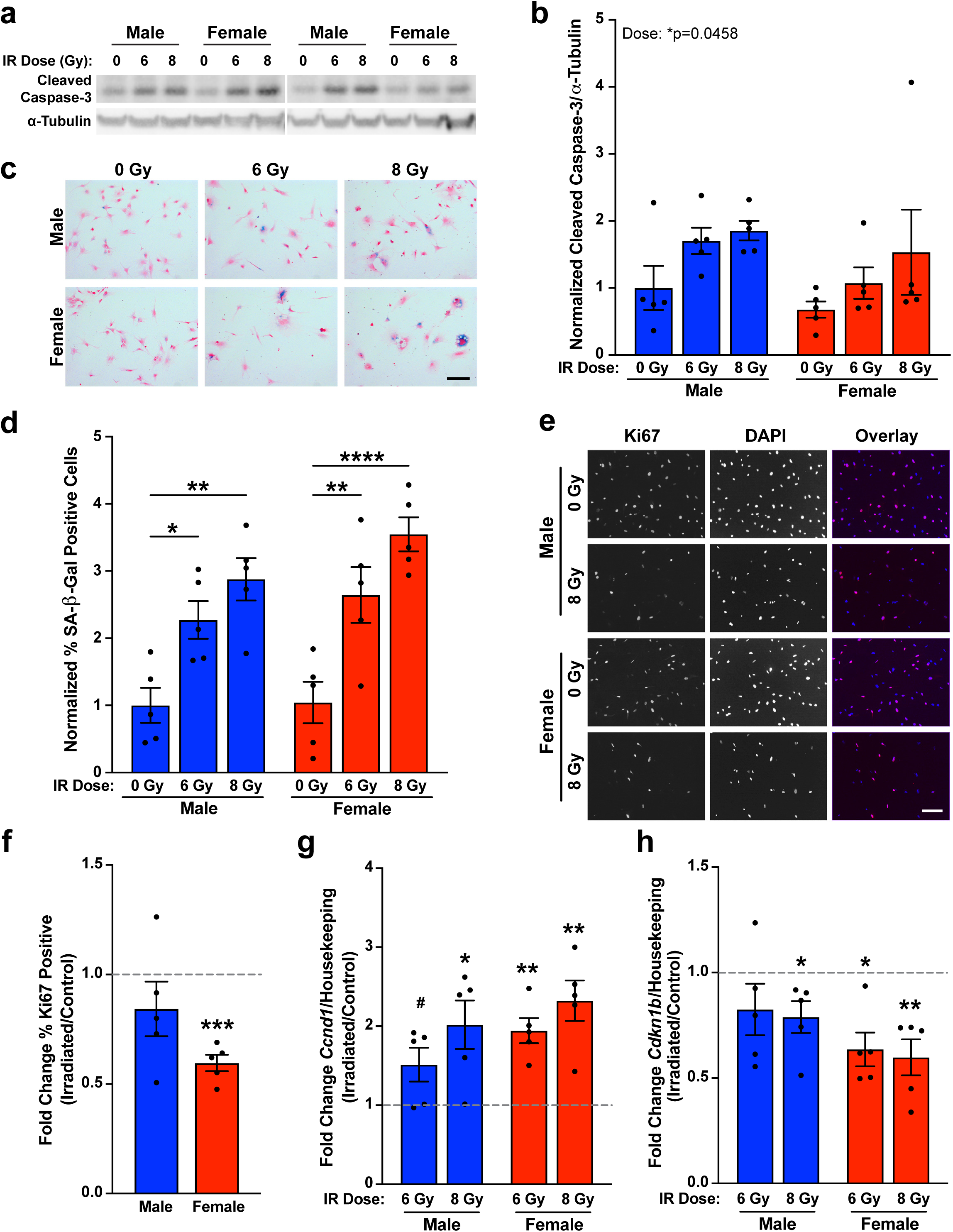
Senescence is a major component of the radiation response in mouse GBM model astrocytes. **a** Western blot images of cleaved caspase-3 5 days after irradiation with 0, 6, or 8 Gy. Representative bands from two *Nf1-/- DNp53* cell lines (male and female) are shown, demonstrating a range of responses. **b** Quantification of cleaved caspase-3 levels measured by western blot. Values were normalized to the corresponding Male 0 Gy condition of the same cell line, which was arbitrarily set at 1. Two-way ANOVA: Dose p=0.0458, Sex p=0.124, Interaction p=0.8633. Data are means +/-SEM (n=5/sex/dose). **c** Example images of *Nf1-/- DNp53* astrocytes stained for SA-β-gal (blue) 5 days after irradiation with 0, 6, or 8 Gy, followed by counterstaining with nuclear fast red (pink). Scale bar, 150 μm. **d** Quantification of the percentage of SA-β-gal positive cells. Values were normalized to the corresponding Male 0 Gy condition of the same cell line, which was arbitrarily set at 1. Two-way ANOVA: Dose p<0.0001, Sex p=0.1671, Interaction p=0.6071. *p<0.05, **p<0.01, ****p<0.0001 as indicated by bracket. Data are means +/-SEM (n=5/sex/dose). **e** Example immunofluorescence images of *Nf1-/- DNp53* astrocytes stained for the cell proliferation marker Ki67 5 days after irradiation with 0 or 8 Gy. Nuclei were counterstained with DAPI. Scale bar, 150 μm. **f** Fold change in the percentage of Ki67 positive cells 5 days after irradiation with 0 or 8 Gy. ***p<0.001 vs 1.0, one sample t-test. Data are means +/-SEM (n=5/sex). **g** Fold change in expression of *Ccnd1* (Cyclin D1) mRNA 5 days after irradiation with 0, 6, or 8 Gy, measured by qPCR. ^#^p=0.0747, *p<0.05, **p<0.01 vs 1.0, one sample t-test. Data are means +/-SEM (n=5/sex/dose). **h** Fold change in expression of *Cdkn1b* (p27) mRNA 5 days after irradiation with 0, 6, or 8 Gy, measured by qPCR. *p<0.05, **p<0.01 vs 1.0, one sample t-test. Data are means +/-SEM (n=5/sex/dose).

**Fig. 4:**
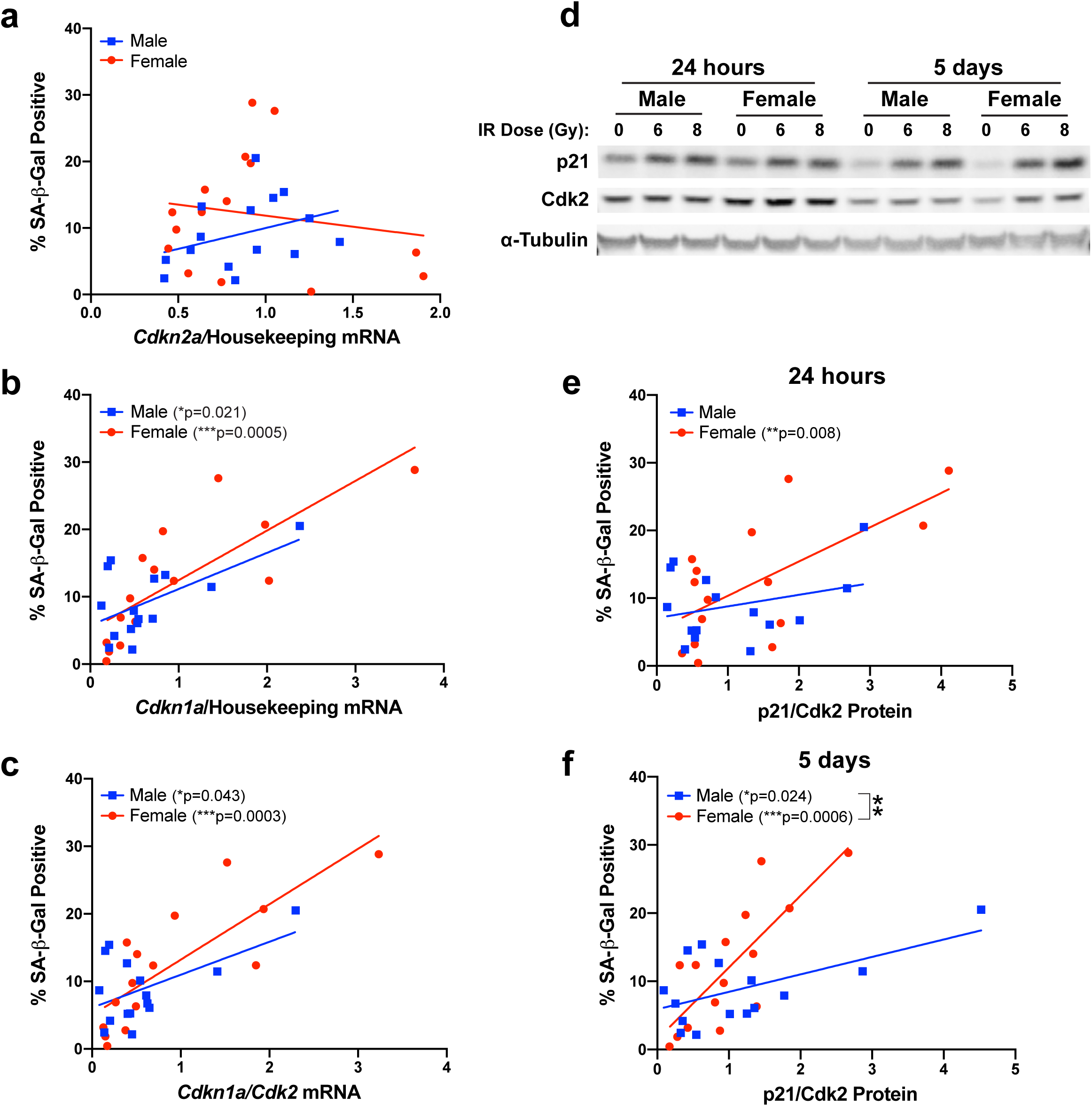
Expression of p21 differentially correlates with SA-β-gal positivity in male and female mouse GBM model astrocytes. **a** Correlation between *Cdkn2a* (p16) mRNA levels at 24 hours after irradiation with 0, 6, or 9 Gy and the percentage of SA-β-gal positive cells at 5 days after irradiation in male and female *Nf1-/- DNp53* astrocytes (n=15/sex – 5 lines, 3 doses). **b** Correlation between *Cdkn1a* (p21) mRNA levels at 24 hours after irradiation with 0, 6, or 9 Gy and the percentage of SA-β-gal positive cells at 5 days after irradiation in male and female *Nf1-/- DNp53* astrocytes. Male correlation: r=0.59, p=0.0212; Female correlation: r=0.79, p=0.0005 (n=15/sex – 5 lines, 3 doses). **c** Correlation between the *Cdkn1a/Cdk2* mRNA ratio at 24 hours after irradiation with 0, 6, or 9 Gy and the percentage of SA-β-gal positive cells at 5 days after irradiation in male and female *Nf1-/- DNp53* astrocytes. Male correlation: r=0.53, p=0.0427; Female correlation: r=0.80, p=0.0003 (n=15/sex – 5 lines, 3 doses). **d** Representative western blot images of p21 and Cdk2 24 hours and 5 days after irradiation with 0, 6, or 8 Gy in one male and one female *Nf1-/- DNp53* cell line. **e** Correlation between the p21/Cdk2 protein ratio at 24 hours after irradiation with 0, 6 or 8 Gy and the percentage of SA-β-gal positive cells at 5 days after irradiation in male and female *Nf1-/- DNp53* astrocytes. Female correlation: r=0.65, p=0.0083 (n=15/sex – 5 lines, 3 doses). **f** Correlation between the p21/Cdk2 protein ratio 5 days after irradiation with 0, 6, or 8 Gy and the percentage of SA-β-gal positive cells at 5 days after irradiation in male and female *Nf1-/- DNp53* astrocytes. Male correlation: r=0.58, p=0.0244; Female correlation: r=0.78, p=0.0006. Male slope vs Female slope: p=0.0026 – indicated by bracket (n=15/sex – 5 lines, 3 doses).

**Fig. 5:**
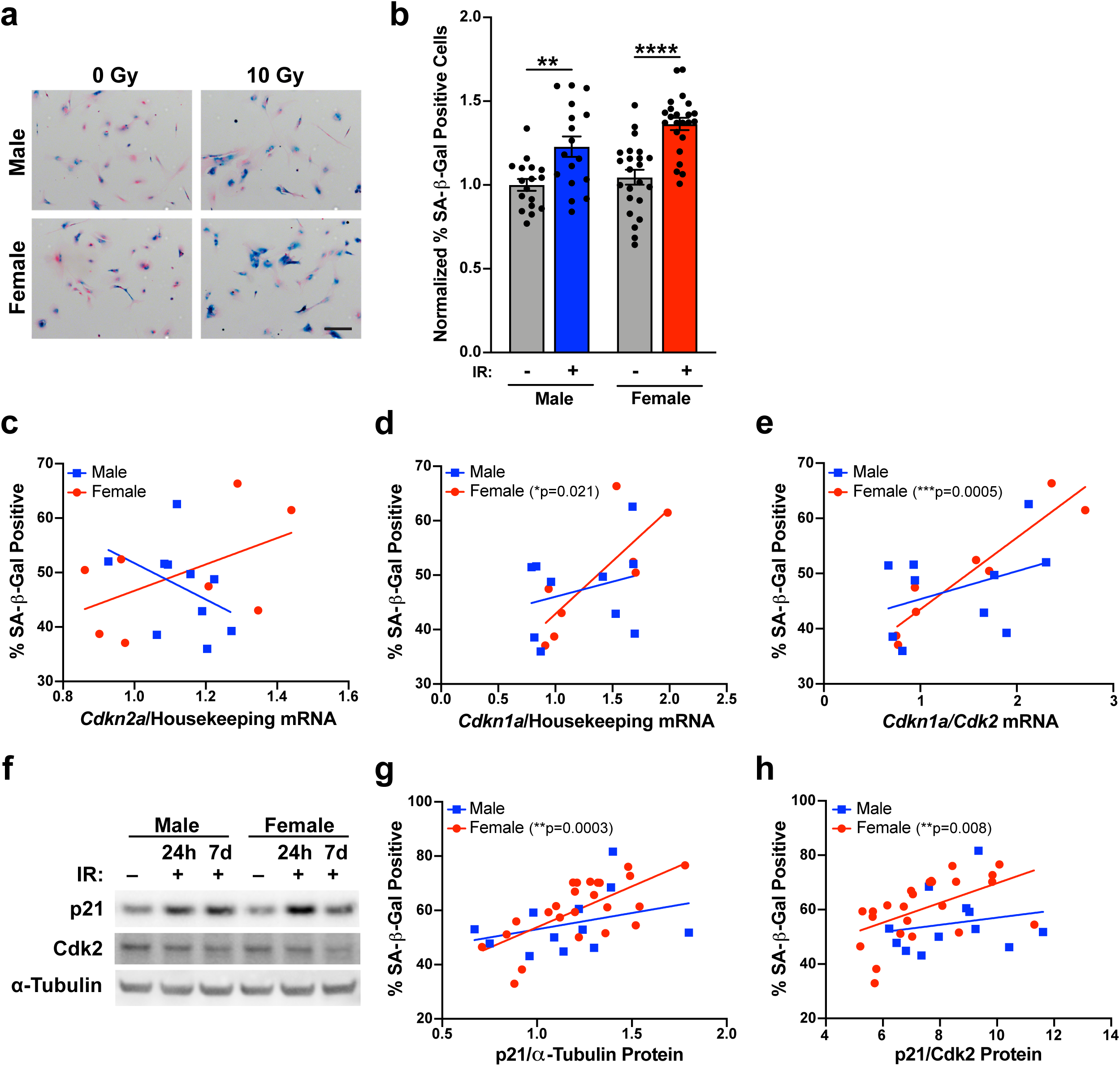
Wildtype mouse astrocytes exhibit sex differences in the relationship between p21 and SA-β-gal positivity. **a** Example images of wildtype mouse astrocytes stained for SA-β-gal seven days after irradiation with 0 or 10 Gy, followed by counterstaining with nuclear fast red. Scale bar, 150 μm. **b** Quantification of the percentage of SA-β-gal positive cells. Results are pooled from 3 separate experiments, representing a total of 17 male and 23 female pups across 7 litters. Values were normalized to the Male 0 Gy condition of the same experiment, which was arbitrarily set at 1. Two-way ANOVA: Dose p<0.0001, Sex p=0.0455, Interaction p=0.3676. **p<0.01, ****p<0.0001 as indicated by bracket. Data are means +/-SEM (n=17-23/sex/condition). **c** Correlation between *Cdkn2a* (p16) mRNA levels at 24 hours after irradiation with 0 or 10 Gy and the percentage of SA-β-gal positive cells at 7 days after irradiation in male and female wildtype astrocytes (n=10 male – 5 pups, 2 doses/8 female – 4 pups, 2 doses). **d** Correlation between *Cdkn1a* (p21) mRNA levels at 24 hours after irradiation with 0 or 10 Gy and the percentage of SA-β-gal positive cells at 7 days after irradiation in male and female wildtype astrocytes. Female correlation: r=0.79, p=0.0207 (n=10 male – 5 pups, 2 doses/8 female – 4 pups, 2 doses). **e** Correlation between the *Cdkn1a/Cdk2* mRNA ratio at 24 hours after irradiation with 0 or 10 Gy and the percentage of SA-β-gal positive cells at 7 days after irradiation in male and female wildtype astrocytes. Female correlation: r=0.94 p=0.0005 (n=10 male – 5 pups, 2 doses/8 female – 4 pups, 2 doses). **f** Representative western blot images of p21 and Cdk2 in non-irradiated and irradiated male and female wildtype astrocytes at 24 hours, and irradiated male and female wildtype astrocytes at 7 days. **g** Correlation between p21 protein levels at 24 hours after irradiation with 0 or 10 Gy and the percentage of SA-β-gal positive cells at 7 days after irradiation in male and female wildtype astrocytes. Female correlation: r=0.67, p=0.0003 (n=12 male – 6 pups, 2 doses/24 female – 12 pups, 2 doses). **h** Correlation between the p21/Cdk2 protein ratio at 24 hours after irradiation with 0 or 10 Gy and the percentage of SA-β-gal positive cells at 7 days after irradiation in male and female wildtype astrocytes. Female correlation: r=0.53, p=0.0081 (n=12 male – 6 pups, 2 doses/24 female – 12 pups, 2 doses).

**Fig. 6:**
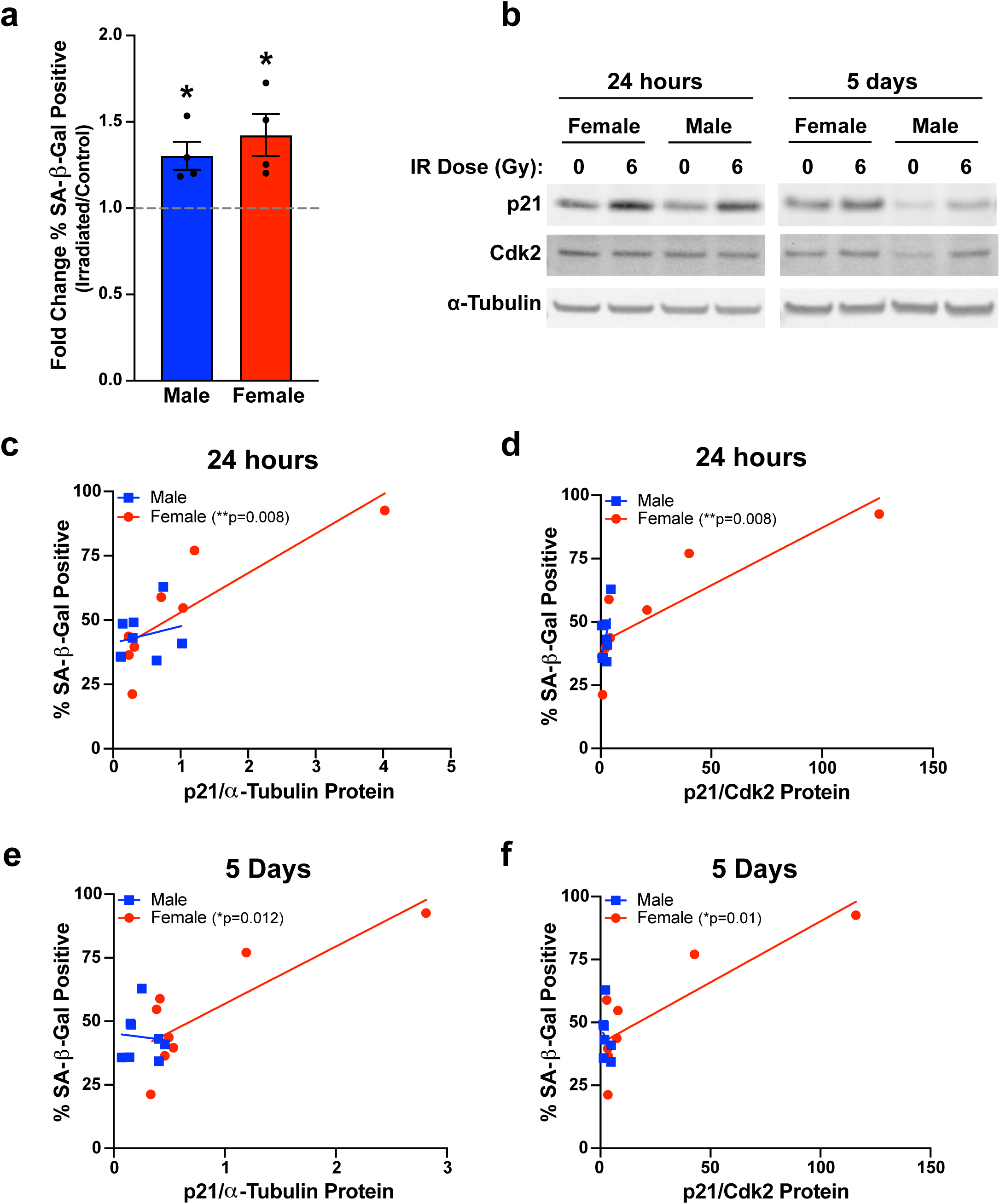
Primary human GBM lines exhibit sex differences in the relationship between p21 and SA-β-gal positivity. **a** Fold change in the percentage of SA-β-gal positive cells 5 days after irradiation with 0 or 6 Gy in 4 male and 4 female primary human GBM lines. *p<0.05 vs 1.0, one sample t-test. Data are means +/-SEM (n=4/sex). **b** Representative western blot images of p21 and Cdk2 24 hours and 5 days after irradiation with 0 or 6 Gy in male and female human GBM lines. **c** Correlation between p21 protein levels at 24 hours after irradiation with 0 or 6 Gy and the percentage of SA-β-gal positive cells at 5 days after irradiation in male and female human GBM lines. Female correlation: r=0.85, p=0.008 (n=8/sex – 4 lines, 2 doses). **d** Correlation between the p21/Cdk2 protein ratio at 24 hours after irradiation with 0 or 6 Gy and the percentage of SA-β-gal positive cells at 5 days after irradiation in male and female human GBM lines. Female correlation: r=0.85, p=0.0081 (n=8/sex – 4 lines, 2 doses). **e** Correlation between p21 protein levels at 5 days after irradiation with 0 or 6 Gy and the percentage of SA-β-gal positive cells at 5 days after irradiation in male and female human GBM lines. Female correlation: r=0.83, p=0.0115 (n=8/sex – 4 lines, 2 doses). **f** Correlation between the p21/Cdk2 protein ratio at 5 days after irradiation with 0 or 6 Gy and the percentage of SA-β-gal positive cells at 5 days after irradiation in male and female human GBM lines. Female correlation: r=0.84, p=0.0099 (n=8/sex – 4 lines, 2 doses).

**Fig. 7:**
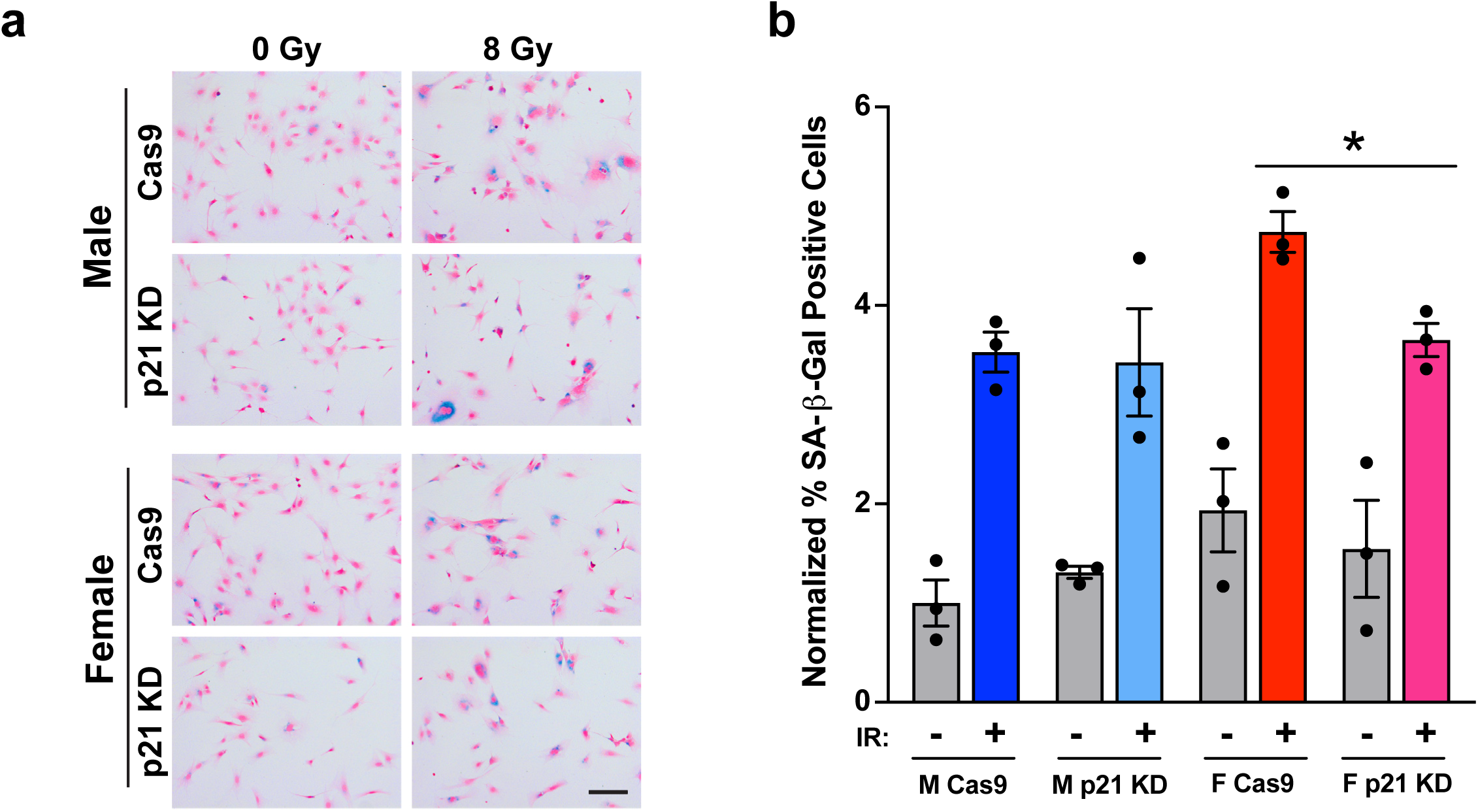
p21 knockdown decreases senescence in irradiated female GBM model astrocytes. **a** Example images of *Nf1-/- DNp53* Cas9 control and p21 knockdown male and female astrocytes stained for SA-β-gal 5 days after irradiation with 0 or 8 Gy, followed by counterstaining with nuclear fast red. Scale bar, 150 μm. **b** Quantification of the percentage of SA-β-gal positive cells. Results are from 3 separate experiments. Values were normalized to the Male Cas9 0 Gy condition, which was arbitrarily set at 1. *p<0.05 as indicated by bracket, two-tailed t-test. Data are means +/-SEM (n=3/group).

### Female mouse GBM model astrocytes are more sensitive to radiation treatment

In order to better understand the mechanisms underlying radiation response in male and female GBM, we utilized an *in vitro* mouse model. This model consists of murine astrocytes with loss of function of the tumor suppressors Nf1 and p53 (*Nf1-/- DNp53*)^13^. When implanted intracranially, both male and female cells form tumors that histologically resemble high grade gliomas, although the frequency of tumor formation differs by cell sex^13^. Additionally, this model displays sex differences in gene expression that are concordant with sex differences in human GBM patient gene expression^14^. We performed irradiation dose response curves with male and female *Nf1-/- DNp53* astrocytes, and found a significant sex difference (Fig. 2a). Female cells were more sensitive to irradiation, a finding consistent with results from human GBM that suggest female GBM patients are more responsive to standard of care therapy (radiation and chemotherapy)^7^.

To further assess sex differences in radiation sensitivity, we irradiated male and female *Nf1-/- DNp53* astrocytes with 0, 3, or 9 Gy, then used live cell imaging to track cell growth for 3 days (Fig. 2b). As previously reported^13^, in the absence of radiation treatment male cells grew faster than female cells. While both male and female cells showed impairment in growth after irradiation with 9 Gy, only female cells showed a decline in growth after treatment at the lower dose of 3 Gy (Fig. 2b). This suggests that female GBM model astrocytes are more sensitive to low dose irradiation. We next used the colony formation assay to assess clonogenic survival following radiation treatment. Female *Nf1-/- DNp53* astrocytes had a greater decrease in colony counts with radiation treatment than male *Nf1-/- DNp53* astrocytes (Fig. 2c), further supporting the idea that female GBM model cells are more sensitive to irradiation.

Finally, we assessed DNA damage resolution by irradiating male and female *Nf1-/- DNp53* astrocytes with 3 Gy, then staining for γH2AX, a marker of double strand breaks (Fig. 2d). We quantified the percent of cells with greater than 10 γH2AX foci at 1, 6, 24, and 48 hours after irradiation (Fig. 2e). At 1 hour after treatment, essentially 100% of cells were positive in both males and females. However, over time the percent of positive cells declined more rapidly in males than in females. At 48 hours, males had returned to baseline levels of DNA damage, while the percent of positive cells remained elevated in females (Fig. 2e). This suggests that radiation treatment results in more persistent DNA damage in female cells. Together these results indicate that female GBM model cells are more sensitive to irradiation than male GBM model cells, a finding which may provide insight into why female GBM patients have a survival advantage.

### Senescence is a major component of the response to irradiation in mouse GBM model astrocytes

Cell response to DNA damage can vary. Cells may undergo: 1) transient cell cycle arrest and repair of DNA damage, 2) apoptosis, or 3) permanent cell cycle arrest in the form of senescence^15^. We focused on the two mechanisms that result in durable therapeutic responses to cancer therapy – apoptosis and senescence. To assess apoptosis, we measured levels of cleaved caspase-3 five days after irradiation (Fig. 3a, b). Overall, we observed a small, but significant treatment-induced increase in cleaved caspase-3 by Two-Way ANOVA. There was no effect of sex on response (Fig. 3b). Measurement of cleaved PARP levels did not confirm the treatment effect (Supplementary Fig. 2a, b), nor did we see any significant change in either apoptosis marker at 24 hours after irradiation (Supplementary Fig. 2a, c, d). Thus, apoptosis does not appear to be the primary mechanism responsible for the decrease in cell number following irradiation.

To measure senescence, we stained cells for senescence-associated β-galactosidase (SA-β-gal), the most widely used biomarker of senescence^11, 16, 17^, five days after irradiation, then quantified the percentage of SA-β-gal positive cells (Fig. 3c, d). In response to irradiation, there was a clear, dose dependent increase in the percentage of senescent cells in both male and female *Nf1-/- DNp53* astrocytes (Fig. 3d). Consistent with an increase in senescent cells, we observed a change in cell morphology after irradiation^11, 16^, with a greater percentage of cells appearing enlarged and irregular in shape (Supplementary Fig. 3). In addition, when we stained for the cell proliferation marker Ki67, we saw a decrease in the percent of positive cells after irradiation in both sexes, although this decrease was more variable in the male cell lines (Fig. 3e, f). Together, these findings are consistent with an increase in senescence. To further confirm the senescence response, we assessed expression levels of two genes previously identified as part of an ionizing radiation induced senescence (IRIS) signature that was shared across multiple cell types and time points^18^. In agreement with this previous report, we observed an increase in expression of *Ccnd1* (Cyclin D1) (Fig. 3g) and a decrease in expression of *Cdkn1b* (p27) (Fig. 3h) 5 days after irradiation in both male and female *Nf1-/- DNp53* astrocytes, providing further support that radiation induced senescence is occurring in these cells. These findings suggest that senescence is a central component of the response to irradiation in *Nf1-/- DNp53* astrocytes, and that this, rather than apoptosis, primarily drives the decrease in cell growth after radiation.

### Expression of p21 24 hours after irradiation correlates with the senescence response observed at 5 days

Two critical pathways for the induction of cellular senescence are the p16^INK4A^/Rb and p53/p21^WAF1/Cip1^ pathways^10, 11^. While the *Nf1-/- DNp53* astrocytes express a dominant negative p53, and thus lack p53 function, they retain the ability to upregulate p21 in response to DNA damage^14^, presumably through p53-independent mechanisms. In order to better understand which of these two pathways may underlie induction of senescence following irradiation, we used qPCR to measure expression levels of *Cdkn2a* (p16) and *Cdkn1a* (p21) 24 hours after treatment of cells with 0, 6, or 9 Gy (for nomenclature clarification, we will use the gene name (*Cdkn2a, Cdkn1a*) when referring to mRNA expression levels, and the protein name (p16, p21) when referring to protein expression levels). We then correlated these values with the percentage of cells that were SA-β-gal positive at 5 days. Because our lab has previously identified sex differences in the regulation of both p16 and p21 in *Nf1-/- DNp53* astrocytes^14^, we looked at the relationship between these two measures and senescence in males and females separately. *Cdkn2a* levels 24 hours after irradiation showed no significant correlation with the senescence response at 5 days in either males or females (Fig. 4a, Supplementary Table 3). In contrast, levels of *Cdkn1a* significantly correlated with the percent of SA-β-gal positive cells in both males (r=0.59) and females (r=0.79) (Fig. 4b, Supplementary Table 3). This suggests that early upregulation of p21, but not p16, contributes to the senescence response following irradiation in mouse GBM astrocytes. This is consistent with previous research from our lab, which found that in response to treatment with the DNA damaging agent etoposide, levels of p21, but not p16, increased in Nf1-/-DNp53 cells^14^.

Downstream of p21 is the cyclin dependent kinase Cdk2, and it is through inhibition of Cdk2 that p21 primarily acts to maintain Rb in a hypophosphorylated state^19, 20^, a critical step in senescence induction^11^. Additionally, it has been reported that the p21/Cdk2 ratio is the primary determinant of the senescent fate decision in human fibroblasts following irradiation^21^. To assess whether the p21/Cdk2 ratio may similarly influence the induction of senescence in our mouse GBM model cells, we measured levels of *Cdk2* mRNA 24 hours after irradiation and correlated the *Cdkn1a/Cdk2* ratio with the percent of SA-β-gal positive cells at 5 days (Fig. 4c, Supplementary Table 3). We found that the *Cdkn1a/Cdk2* ratio significantly correlated with SA-β-gal positivity in both males (r=0.53) and females (r=0.80).

### Expression of p21 differentially correlates with SA-**β**-gal positivity in male and female mouse GBM model astrocytes

We next sought to determine whether the correlation between p21/Cdk2 ratio and senescence was also observed at the protein level, and whether this relationship was maintained at later timepoints, once senescence was established. To this end, we measured p21 and Cdk2 levels by western blot, 24 hours and 5 days after irradiation with 0, 6, and 8 Gy (Fig. 4d-f). Surprisingly, while p21/Cdk2 protein levels at 24 hours significantly correlated with SA-β-gal positivity in females (r=0.65), this was not the case for males (r=0.27) (Fig. 4e, Supplementary Table 3). Even more striking, while p21/Cdk2 protein levels at 5 days did correlate with the percentage of SA-β-gal positive cells in both males (r=0.58) and females (r=0.78), it was with significantly different slopes (Fig. 4f, Supplementary Table 3). The rate of increase was much faster in females (slope estimate/standard error (SE)=10.53/2.07) than males (slope estimate/SE=2.59/1.18) (slope difference p=0.0026). Thus, the same level of p21/Cdk2 expression at 5 days corresponded to higher levels of senescence in females than in males, as measured by SA-β-gal. This suggests that not only does the p21/Cdk2 ratio play a role in the maintenance of senescence following irradiation – but that there may be a sex difference in sensitivity to p21/Cdk2 levels, and/or that p21 may be playing a greater role in irradiation induced senescence in female GBM cells, than in male GBM cells.

### Wildtype mouse astrocytes exhibit sex differences in the relationship between p21 and SA-**β**-gal

We wondered whether the sex difference in the relationship between p21 and SA-β-gal that we observed in our mouse GBM model represents a fundamental sex difference, present in normal astrocytes, or is unique to transformed cells and the loss of p53 function. To address this question, we isolated astrocytes from the cortex of male and female postnatal day 1 C57Bl6 mouse pups. Astrocytes from each pup were cultured independently and split to allow for corresponding measures of SA-β-gal and collection of RNA or protein. We irradiated these wildtype (WT) astrocytes with 0 or 10 Gy, then collected RNA or protein 24 hours later, and stained for SA-β-gal at 7 days (Fig. 5a). The irradiation dose and time until SA-β-gal measurement were increased to adjust for the much slower division rate of wildtype astrocytes compared to *Nf1-/- DNp53* astrocytes. Quantification of the percentage of SA-β-gal positive cells confirmed a significant increase following irradiation in both male and female wildtype astrocytes (Fig. 5b).

We used qPCR to measure expression levels of *Cdkn2a* (p16) and *Cdkn1a* (p21) mRNA at 24 hours after irradiation, then correlated these measures with the corresponding SA-β-gal percentage. As expected, *Cdkn2a* levels did not significantly correlate with SA-β-gal positivity in either male or female WT astrocytes (Fig. 5c, Supplementary Table 3), confirming that p16 upregulation is not a primary driver of senescence induction in astrocytes following irradiation. In contrast, *Cdkn1a* levels did significantly correlate with the percent of SA-β-gal positive cells at 7 days, however, only in females (r=0.79, r=0.28 for males) (Fig. 5d, Supplementary Table 3). When we looked at the relationship between the *Cdkn1a/Cdk2* ratio and SA-β-gal, we once again found a significant correlation in female WT astrocytes (r=0.94), but a non-significant mild correlation in male WT astrocytes (r=0.39, male vs. female correlation difference p=0.024) (Fig. 5e, Supplementary Table 3). As with the *Nf1-/- DNp53* astrocytes, the female WT astrocytes had a steeper slope for both *Cdkn1a* and *Cdkn1a/Cdk2* vs SA-β-gal compared to males, although this did not reach statistical significance.

To determine whether the sex difference in the relationship between p21 expression and SA-β-gal was maintained at the protein level, we measured p21 and Cdk2 protein levels by western blot (Fig. 5f), then correlated these with the percent of SA-β-gal positive cells at 7 days. Both p21 alone, and the p21/Cdk2 ratio, significantly correlated with senescence in female WT astrocytes (r=0.67 and r=0.53 respectively), but not male WT astrocytes (Fig. 5g, h, Supplementary Table 3). These results suggest that p21 may be playing a greater role in senescence induction in female astrocytes than in male astrocytes following irradiation, and that this is occurring regardless of whether the cells are transformed or not.

### Sex differences in senescence are observed in wildtype astrocytes with repeated *in vitro* passaging

In initial studies to assess the effects of treatment induced senescence in wildtype astrocytes, we observed that as the cells reached higher passages, a baseline sex difference in senescence began to emerge – with females having higher percentages of SA-β-gal positive cells than males in the untreated condition. To assess this more directly, we cultured male and female wildtype astrocytes and stained for SA-β-gal when the cells were at either low (p2 or p3) or high (p5) passage (Supplementary Fig. 4a). At greater than five passages, the wildtype astrocytes showed a dramatic decrease in growth, with the majority of cells adopting an enlarged, flattened, irregular morphology, and were unable to be passaged further.

At p2/3, the ratio of SA-β-gal positive cells (female/male) was approximately 1, indicating equivalent levels of senescent cells in male and female astrocyte cultures (Supplementary Fig. 4b). At p5, this ratio was significantly increased, indicating higher percentages of senescent cells in female cultures (Supplementary Fig. 4b). Thus, female mouse astrocytes undergo senescence more frequently than male mouse astrocytes with repeated passaging. Cell cycle analysis of male and female wildtype astrocytes by flow cytometry found no difference in cell cycle distribution or cell size based on sex (data not shown), suggesting the difference in senescence is not due to baseline differences in cell proliferation rates. Whether it reflects sex differences in the propensity to undergo replicative senescence or in the sensitivity to oxidative damage, resulting from culture at 20% oxygen^22^, remains to be determined. However, this suggests that sex differences may extend to other senescence paradigms and phenotypes, and highlights the need to study these in both sexes separately.

### Primary human GBM lines exhibit sex differences in the relationship between p21 and SA-**β**-gal

We next assessed whether sex differences in the role of p21 in irradiation induced senescence extended to human GBM. For this purpose, we utilized the same primary human GBM lines used to correlate irradiation sensitivity with expression of the MC5 and FC3 gene sets. We irradiated each male and female line with 0 or 6 Gy and stained for SA-β-gal 5 days later. The 6 Gy dose was chosen based on the human GBM irradiation dose response curves; it was above the mean IC_50_ value for both sexes, but lower than the dose at which most lines showed a plateau in response. There was a significant increase in the percent of SA-β-gal positive cells following irradiation in both male and female human GBM lines (Fig. 6a).

Using protein samples collected 24 hours and 5 days after irradiation, we measured levels of p21 and Cdk2 by western blot (Fig. 6b). At 24 hours, both p21 alone (Fig. 6c) and the p21/Cdk2 ratio (Fig. 6d) significantly correlated with the percent of SA-β-gal positive cells at 5 days in female human GBM lines (r=0.85 for both), but only moderately correlated in male human GBM lines (r=0.23 and r=0.54, respectively) (Supplementary Table 3). This same pattern was observed at 5 days, with p21 and p21/Cdk2 significantly correlating with SA-β-gal for female (r=0.83 and r=0.84, respectively) but not male lines (r=-0.09 and r=-0.27, respectively, p21 male vs. female correlation difference p=0.0461, p21/Cdk2 male vs. female correlation difference p=0.0192) (Fig. 6e, f, Supplementary Table 3). Thus, in both mouse and human, transformed and wildtype astrocytes, the relationship between p21 and senescence, as measured by SA-β-gal, differs between males and females.

### Knockdown of p21 decreases senescence in irradiated female mouse GBM model astrocytes

To further test the role of p21 in senescence in irradiated male and female cells, we utilized an *Nf1-/- DNp53* p21 knockdown line previously developed in our lab using CRISPR/Cas9^14^. We confirmed knockdown by irradiating Cas9 and p21 KD cells with 0 or 8 Gy and measuring levels of p21 protein 24 hours later. Both male and female p21 KD lines had a strong reduction in p21 levels – less than 30% of control levels (Supplementary Fig. 5a, b). We then irradiated male and female Cas9 and p21 KD cells with 0 or 8 Gy and stained for SA-β-gal 5 days later (Fig. 7a). The female p21 KD line had a significant reduction in the percent of SA-β-gal positive cells compared to female Cas9 after irradiation, decreasing to levels similar to those of male cells (Fig. 7b). This supports the idea that p21 expression contributes significantly to the levels of senescence after irradiation in females, but not in males. Surprisingly, despite the decrease in SA-β-gal in female p21 KD cells, they still retain a significant senescence response, and in fact decrease only to the levels of senescence in male cells. This suggests that additional pathways, beyond p21, are contributing to senescence after irradiation.

### The Four Core Genotypes mouse model can be used to interrogate the developmental mechanisms underlying sex differences

Multiple mechanisms can potentially underlie an observed sex difference: 1) acute differences in circulating gonadal hormone levels, 2) organizational/epigenetic effects of gonadal hormone exposure *in utero*, and 3) differences in the expression levels of genes on the X and Y chromosomes^23^. The sex difference in the relationship between p21 and SA-β-gal that we observe is present *in vitro,* with male and female cells grown in identical media. Thus, it is unlikely that the differences in p21 are driven by effects of circulating estrogen or testosterone. To investigate whether organizational effects or sex chromosomes are responsible, we utilized a transgenic mouse model known as the Four Core Genotypes (FCG) model^24^. In the FCG model, the *Sry* gene, which encodes the testis-determining factor, is deleted from the Y chromosome and inserted onto an autosome, allowing for the separation of gonadal and chromosomal sex. Crossing XY^-^/Sry+ males with normal XX females results in mice of four genotypes: XY^-^/Sry+ (XY with testes – normal male), XY^-^/Sry-(XY with ovaries), XX/Sry+ (XX with testes), and XX/Sry-(XX with ovaries - normal female) (Fig. 8a). Mice that inherit the *Sry* gene, regardless of chromosomal sex, develop testes and are exposed to masculinizing levels of gonadal hormones during *in utero* development. Mice that lack the *Sry* gene develop ovaries and display phenotypes associated with a feminized brain^24^. We crossed FCG XY^-^/Sry+ mice with Cas9 expressing female (XX) mice and isolated astrocytes from the postnatal day 1 pups. We then used CRISPR/Cas9 to delete Nf1 and p53 from these astrocytes, mimicking our *Nf1-/- DNp53* GBM model (Supplementary Fig. 6).

**Fig. 8:**
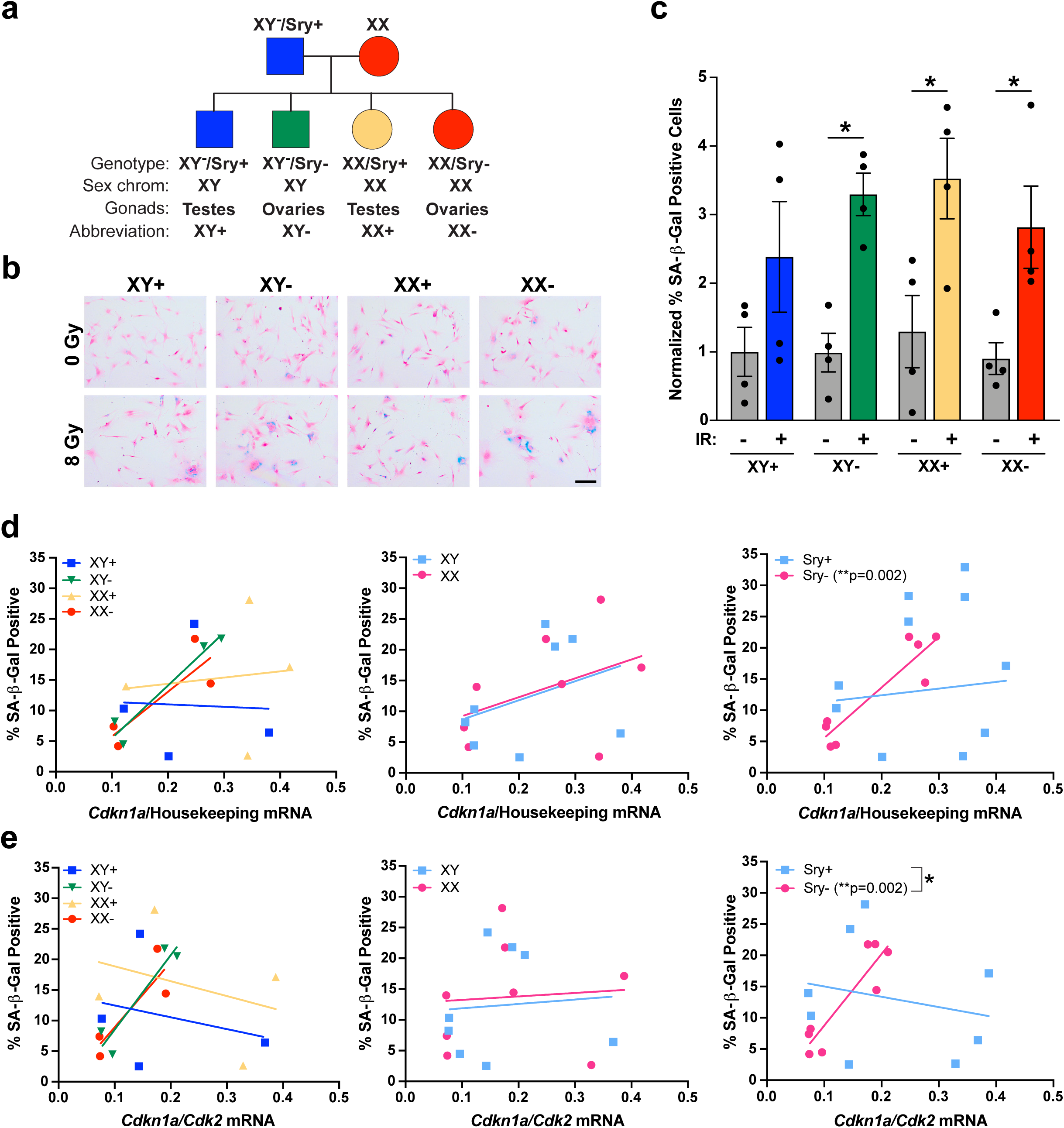
Gonadal sex patterns the relationship between p21 and SA-β-gal positivity. **a** Diagram of the four genotypes resulting from the FCG mouse model. **b** Example images of FCG GBM model astrocytes stained for SA-β-gal 5 days after irradiation with 0 or 8 Gy, followed by counterstaining with nuclear fast red. Scale bar, 150 μm. **c** Quantification of the percentage of SA-β-gal positive cells. Results are from two cell lines (each consisting of a corresponding XY+, XY-, XX+, and XX-line generated at the same time) tested in 2 separate experiments. Values were normalized to the XY+ 0 Gy condition, which was arbitrarily set at 1. *p<0.05 as indicated by bracket. Data are means +/-SEM (n=4/genotype/condition). **d** Correlation between *Cdkn1a* (p21) mRNA levels at 24 hours after irradiation with 0 or 8 Gy and the percentage of SA-β-gal positive cells at 5 days after irradiation in FCG GBM model astrocytes, showing each genotype separately (left), as well as grouped by sex chromosomes (center) or gonadal sex (Sry status) (right). Sry- correlation: r=0.91, p=0.0015 (n=8/group – 4 lines, 2 doses). **e** Correlation between the *Cdkn1a/Cdk2* mRNA ratio at 24 hours after irradiation with 0 or 8 Gy and the percentage of SA-β-gal positive cells at 5 days after irradiation in FCG GBM model astrocytes, showing each genotype separately (left), as well as grouped by sex chromosomes (center) or gonadal sex (Sry status) (right). Sry- correlation: r=0.91, p=0.002. Sry+ slope vs Sry- slope: p=0.0262 – indicated by bracket (n=8/group – 4 lines, 2 doses).

### Gonadal sex patterns the relationship between p21 and SA-**β**-gal positivity

Using our newly developed FCG GBM model, we irradiated the cells with 0 or 8 Gy and then stained for SA-β-gal 5 days later (Fig. 8b). The percentage of SA-β-gal positive cells increased following irradiation in all four genotypes (Fig. 8c), although due to high variability this did not reach significance in the XY+ genotype. Using qPCR, we measured levels of *Cdkn1a* (p21) mRNA expression 24 hours after irradiation and correlated this with the percent of SA-β-gal positive cells at 5 days (Fig. 8d, left, Supplementary Table 3). To better evaluate whether sex chromosomes or gonadal sex determines the relationship between p21 and SA-β-gal, we grouped the genotypes based on these factors and then compared XY vs XX (Fig. 8d, center, Supplementary Table 3) and Sry+ vs Sry- (Fig. 8d, right, Supplementary Table 3). When the genotypes were grouped based on sex chromosomes, *Cdkn1a* did not significantly correlate with SA-β-gal in either XY or XX FCG GBM model cells. However, when the genotypes were grouped based on gonadal sex, *Cdkn1a* expression significantly correlated with SA-β-gal positivity in Sry- cells (r=0.91), but not Sry+ cells (r=0.13, Sry+ vs. Sry- correlation difference p=0.03).

We next measured *Cdk2* levels by qPCR and correlated the *Cdkn1a/Cdk2* ratio with SA-β-gal percentage (Fig. 8e, left, Supplementary Table 3). When we grouped the genotypes based on sex chromosomes, we again observed that the *Cdkn1a/Cdk2* ratio did not correlate with SA-β- gal in either XY or XX cells (Fig. 8e, center, Supplementary Table 3). In contrast, when we grouped the genotypes based on gonadal sex, the *Cdkn1a/Cdk2* ratio significantly correlated with SA-β-gal in Sry- cells (r=0.91), but not Sry+ cells (r=-0.22, Sry+ vs. Sry- correlation difference p=0.01) (Fig. 8e, right, Supplementary Table 3). Additionally, when we compared the slopes of the lines for Sry+ and Sry-, they were significantly different (Sry+ slope estimate/SE=- 1.56/46.98, Sry- slope estimate/SE=131.06/51.73, slope difference p=0.0262). Thus, FCG GBM astrocytes isolated from mice that were gonadally female had a significant relationship between p21 and senescence, as measured by SA-β-gal. This suggests that gonadal sex, and the epigenetic effects of *in utero* gonadal hormones, patterns the role of p21 in senescence induction in response to irradiation.

## Discussion

Glioblastoma remains an incurable disease, with limited treatment options. Surgical resection, followed by radiation and chemotherapy remains the most effective treatment strategy. Better understanding of the mechanisms underlying the therapeutic response to radiation may help enhance treatment efficacy. Here we show that radiation response correlates with different molecular signatures in male and female human GBM lines. Using a mouse model of GBM, we further identify increased sensitivity to radiation in female GBM astrocytes, and determine that senescence is a major component of the radiation response in both sexes. With a correlation-based approach, we identified p21 as a likely mediator of irradiation induced senescence. Strikingly, the relationship between p21 and the senescence marker SA-β-gal significantly differed in male and female cells – a finding that was observed in mouse GBM model astrocytes, mouse wildtype astrocytes, and primary human GBM lines. Female cells had higher levels of senescence, as measured by SA-β-gal, in response to the same levels of p21 – suggesting that p21 plays a more critical role in irradiation induced cellular senescence in female than in male cells. This finding was confirmed by p21 knockdown, which decreased the percentage of senescent cells in female GBM model astrocytes to that of males. Finally, using a novel FCG model of GBM, which enables mechanistic dissection of the biological underpinnings of a sex difference, we determined that the relationship between p21 and irradiation induced cellular senescence is patterned by gonadal sex. This finding will help guide future studies that aim to uncover the pathways regulating the p21-senescence axis and eventually modulate this interaction to improve radiation response in both sexes.

Our results highlight some intriguing directions for future studies. Interestingly, when p21 was knocked down in female GBM model astrocytes, it did not eliminate the senescence response to irradiation, but did abrogate the sex difference in response. This suggests that there are additional pathway(s), beyond p21, contributing to senescence induction after radiation, and that these are shared by males and females. Which signaling pathways, and whether they are always active or are upregulated with p21 loss, remains to be determined. Another avenue of future direction is uncovering how gonadal sex patterns the relationship between p21 and senescence. Our findings suggest that the testosterone surge *in utero* (or lack thereof in females) is responsible for the sex differences in the p21-senescence axis. The hormone surge has been shown to exert long term organizational effects in the brain through epigenetic mechanisms^1, 25^. However, the differences we observe are not explained through simple regulation of accessibility at the p21 locus, since equivalent levels of p21 in males and females correspond to different percentages of SA-β-gal positive cells. Instead, differences in hormone exposure *in utero* may regulate proteins that influence p21 localization or activity through post-translational modifications^26–29^. Future studies will investigate this possibility, with the goal of enhancing the senescence response after irradiation.

We also observed a sex difference in the levels of SA-β-gal cells in male and female wildtype astrocytes with continued passaging *in vitro*. This suggests female cells may be more susceptible to replicative senescence and/or to oxidative stress, since cells were grown at supraphysiological oxygen levels^22^. This has important implications for the fields of aging and neurodegenerative disease. Senescent cells contribute to a wide range of normal and pathological changes with aging^30, 31^, and are increasingly thought to play a role in neurodegenerative disease^32, 33^. Sex differences in p16 and p21 expression in aged mice *in vivo* have been reported, although this study actually found higher levels of these in male mice than female mice^34^. In our study, the same levels of p21 were associated with increased SA-β-gal cells in females, raising the possibility that this difference may not actually translate into increased senescent cells in males. Alternatively, it is possible that the senescent cell phenotype may also differ between male and female cells. The senescence associated secretory phenotype varies based on cell type and senescence induction mechanism^18, 35^, and so could reasonably be influenced by cell sex. This could potentially result in different rates of clearance for male and female senescent cells *in vivo*, or mean that the same levels of senescence have different effects on the surrounding tissue depending on sex.

One of the potential limitations of our study is that we rely on SA-β-gal for our primary measure of senescence. While SA-β-gal is the most widely used marker of senescence, there is debate about its specificity, since it is not required for senescence induction and has been observed in non-senescent cells^16^. In support of the idea that SA-β-gal does correspond to senescence in our study: 1) it increased with exposure to radiation in a dose dependent manner, 2) along with increases in SA-β-gal, we saw more cells with an enlarged, irregular shape and a decrease in cells positive for the proliferation marker Ki67, 3) expression changes consistent with ionizing radiation induced senescence were observed in our cells, and 4) levels of the cell cycle inhibitor p21 also increased. Regardless of whether SA-β-gal is a measure of true senescence, our findings still have important implications for the field. When interpreting studies that use p21 as a marker of senescence, it may be necessary to consider the sex of the subjects, and whether the same pathway could have differences in downstream outcomes between male and female cells.

Finally, our results highlight the utility of correlation analyses when studying sex differences. Most phenotypes are not truly sexually dimorphic, but represent a spectrum of values in males and females. While the mean values differ, and the two ends of the spectrum are populated primarily by individuals of one sex or another, there is considerable overlap. In addition, the variability between individuals of a single sex may obscure the differences between the two sexes, requiring a large sample size to detect the difference. With our approach, we discovered a difference in the relationship between p21 and SA-β-gal in males and females that was not apparent when we simply compared mean p21 levels alone. This could offer a powerful investigation strategy that takes advantage of individual variability rather than viewing it as a liability.

In summary, we have uncovered a novel sex difference in the relationship between p21 and irradiation induced cellular senescence that has potential implications for the fields of cancer research, neuroscience, and aging. Better understanding of this mechanism could identify novel approaches to improve response to cancer therapy.

## Methods

### Primary human GBM lines

Primary human GBM lines were kindly provided by Dr. Albert H. Kim. Specimens for culture were obtained prospectively at the time of surgery and cell lines were established as described^36^. Consent was obtained in accordance with a Washington University Institutional Review Board (IRB) approved Human Studies Protocol. Patient charts were reviewed for the molecular characteristics of the tumors/GBM lines (Supplementary Table 1). Available information varied between tumors; where no information was available, that cell in the table was left blank. Human GBM lines were grown on laminin (Sigma) coated Primaria plates in RHB-A media (Takara) supplemented with growth factors EGF (Sigma) and bFGF (Millipore) at 50 ng/ml. Media was replaced with half fresh media every 2-3 days. For passaging, cells were detached with Accutase (Sigma) and split 1:2 to 1:4 depending on cell line. One female human line (B51) was excluded from the SA-β-gal studies, since these cells did not attach to glass coverslips, even when coverslips were coated with laminin.

### Mouse wildtype astrocytes

All animals were used in accordance with an Animal Studies Protocol approved by the Animal Studies Committee of the Washington University School of Medicine, per the recommendations of the Guide for the Care and Use of Laboratory Animals (NIH). Mouse wildtype astrocytes were isolated from the cortex of postnatal day 1 C57BL/6J pups as described^13^. Briefly, the cortices were dissected from the rest of the brain in cold HBSS and the meninges were removed. Isolated cortices were incubated in 500 μl 0.05% Trypsin-EDTA + 0.02 mg/mL DNase I (Roche) for 15 minutes at 37°C. Trypsin was neutralized by the addition of 750 μl media (DMEM/F12 + 10% FBS + 1% penicillin-streptomycin), and tissue was triturated 10-15 times with a p1000 pipet, then filtered through a 40 μm cell strainer. Cell suspension was spun down, and the pellet was resuspended and plated in a poly-L-lysine (ScienCell) coated T25 flask. Cells were grown until confluent (approximately 7 days). To obtain a pure population of astrocytes, confluent flasks were shaken at 225 rpm overnight at 37°C. After shaking, the media was aspirated, removing any floating cells, and flasks were rinsed once with sterile PBS. Remaining attached cells, representing the astrocyte population, were trypsinized and replated in Primaria T75 flasks (Corning). Cells were then expanded for downstream analyses. Prior to experimental treatment, cells were switched to media without penicillin/streptomycin (DMEM/F12 + 10% FBS). For the irradiation studies, all data points represent astrocyte cultures from individual pups. The sex of each culture was determined by genotyping the corresponding pup tail DNA for the X and Y chromosome paralogs *Kdm5c* (*Jarid1c*) and *Kdm5d* (*Jarid1d*) (Forward: CTG AAG CTT TTG GCT TTG AG, Reverse: CCA CTG CCA AAT TCT TTG G)^37^. All irradiation studies were carried out on astrocytes that were passage 2 or 3. For the studies of low and high passage astrocytes, cultures from multiple pups were pooled by sex to generate male and female cultures.

### Mouse GBM model astrocytes

*Nf1-/- DNp53* astrocytes were generated as described^13^. Briefly, astrocytes were isolated from the cortex of postnatal day 1 *Nf1^flox/flox^;GFAP-Cre* mouse pups. Sex was determined by genotyping pup tail DNA for the X and Y chromosome paralogs *Kdm5c* (*Jarid1c*) and *Kdm5d* (*Jarid1d*)^37^. Astrocytes from at least three male and three female pups were then pooled by sex. Male and female *Nf1-/-* astrocyte cultures were infected with a retrovirus encoding EGFP and a flag-tagged dominant-negative p53 (DNp53), consisting of amino acids 1-14 of the transactivation domain, followed by amino acids 309-393. GFP positive-DNp53 expressing cells were selected for by fluorescence-activated cell sorting (FACS). *Nf1-/- DNp53* cells were cultured in DMEM/F12 supplemented with 10% FBS and 1% penicillin-streptomycin. Five individually generated Nf1-/-DNp53 cell lines, each consisting of a paired male and female line, were used for the experiments in this study.

### p21 knockdown lines

The male and female *Nf1-/- DNp53* Cas9 control and p21 knockdown lines were previously generated and published by our lab^14^. Briefly, *Nf1-/- DNp53* astrocytes were infected with the lentiviral Cas9/sgRNA-p21 all-in-one construct (p21 sgRNA sequence: ACTTCGTCTGGGAGCGCGTT). Cas9 control lines were generated by infecting with the lentiviral Cas9 vector without any guide RNA. Following infection, cell lines were selected by puromycin treatment (2.5 μg/ml) for 1-2 weeks. Surviving cells were expanded and p21 knockdown was confirmed by western blot.

### FCG GBM model astrocytes

To generate the FCG-GBM model, Four Core Genotypes XY^-^/Sry+ male mice^24, 38–40^ (Jackson Laboratory #010905) were bred with constitutive Cas9 expressing^41^ female mice (Jackson Laboratory #026179), and astrocytes were isolated from the cortex of postnatal day 1 FCG-Cas9 mouse pups as described^13^. Pup tail DNA was genotyped for Sry and the Y chromosome following the protocol available from Jackson Laboratory (Sry Forward: AGC CCT ACA GCC ACA TGA TA, Sry Reverse: GTC TTG CCT GTA TGT GAT GG, Y Chrom. Forward: CTG GAG CTC TAC AGT GAT GA, Y Chrom. Reverse: CAG TTA CCA ATC AAC ACA TCA C, Internal Control Forward: CAA ATG TTG CTT GTC TGG TG, Internal Control Reverse: GTC AGT CGA GTG CAC AGT TT). Astrocytes from at least three pups from each of the four genotypes were pooled to generate XY^-^/Sry+ (XY+), XY^-^/Sry-(XY-), XX/Sry+ (XX+), and XX/Sry-(XX-) cultures. FCG-Cas9 astrocytes at passage 4 were transfected with a modified pX330 vector with dual sgRNAs targeting Nf1 and p53 (Nf1 sgRNA sequence: GCAGATGAGCCGCCACATCGA, p53 sgRNA sequence: CCTCGAGCTCCCTCTGAGCC). The pX330-Nf1-p53 vector was kindly provided by Dr. Kwanha Yu and Dr. Benjamin Deneen (Baylor College of Medicine). FCG-Cas9 astrocytes that received the pX330-Nf1-p53 plasmid were immortalized and had a significant growth advantage. Since wildtype astrocytes stop dividing after passage 5-6, growth advantage was used to select for astrocytes with successful CRISPR mutation/deletion of Nf1 and p53. FCG-Cas9 Nf1/p53 CRISPR astrocytes were passaged at least 3 times, then protein was collected for confirmation of Nf1 and p53 knockdown by western blot.

### Irradiation

For irradiation experiments, cells were irradiated using an RS 2000 X-ray irradiator (Rad Source Technologies) unless otherwise specified in the specific methods section. Radiation was delivered at a dose rate of ∼1.8 Gy/min with 160 kVp X-rays. Control cells were transported to the irradiator and sat on the bench for the same length of time as cells were in the irradiator.

### Human GBM irradiation dose response curves

Human GBM cells were lifted with Accutase, counted, and 20,000 cells per well were plated in laminin (Sigma) coated 24 well plates. Cells were plated in triplicate wells for each irradiation dose. Cells were irradiated 24 hours later with 0, 2, 4, 6, 8, or 10 Gy, then allowed to grow for 4 days after irradiation, with fresh media added to the wells 2 days after irradiation. On day 4, media was aspirated off, and 200 μl of Accutase was added to each well. Plates were incubated for 3 min in a 37°C incubator, then wells were rinsed with Accutase to detach all cells, and cell suspension was transferred to 1.7 ml tubes on ice. Cell suspension was diluted 1:1 with Trypan blue, and cells were counted with a hemocytometer. Three technical replicates were counted per dose. Percentage of cell number was calculated by dividing the counts for each dose by the average cell number for the 0 Gy condition. For all cell lines except B5, two separate dose curves were performed, and the 6 technical replicates were combined to get the final irradiation dose response curve. Due to slow growth rate, B5 cell number was limited, and only one dose curve, with 3 technical replicates, was performed.

### Mouse GBM irradiation dose response curves

Irradiation dose response curves for mouse GBM model astrocytes were performed using the sulforhodamine B assay as described, with minor modifications^42^. Briefly, *Nf1-/- DNp53* astrocytes were trypsinized, counted, and 1000 cells per well in 200 μl media were plated in 96-well white-walled clear-bottom plates (Corning). 6 technical replicate wells were plated for each irradiation dose. A standard curve for each cell line and sex was plated at the same time by performing a 2-fold serial dilution starting at 100,000 cells per well, and diluting 6 times, plating 0 cells in the final row. Standard curve wells were plated in triplicate for each cell line. 6 hours after plating, standard curves were fixed by the addition of 100 μl ice cold 5% trichloroacetic acid and incubation at 4°C for 1 hour. Standard curves plates were then washed 4 times with tap water and allowed to dry overnight. 24 hours after plating, cells were irradiated with 0, 2, 3, 4, 6, 8, or 9 Gy, then allowed to grow for 4 days. On day 4, plates were fixed with 5% trichloroacetic acid, then washed and dried. After drying, plates were stained by adding 100 μl of 0.057% sulforhodamine B (SRB) diluted in 1% acetic acid to each well and incubating for 30 minutes at room temperature. Plates were then washed 3 times with 1% acetic acid and allowed to dry overnight. SRB was solubilized by adding 100 μl 10 mM Tris-Base (pH 10.5) to each well, then incubating for 1-2 hours at room temperature on a shaker. Absorbance was measured at 510 nm using an Infinite 200 PRO microplate reader (Tecan). Standard curve absorbance values were graphed using GraphPad Prism software, and a linear standard curve was fit. The standard curve for each cell line was used to interpolate the cell number for experimental wells. Percentage of cell number was calculated by dividing the cell counts for each irradiated well by the average cell number for the 0 Gy condition. Technical replicates were averaged to derive a single value for each dose per cell line and sex, and results from each of the 5 *Nf1-/- DNp53* cell lines were combined to generate the final irradiation dose response curves for male and female mouse GBM model astrocytes.

### Cell growth assays via live cell imaging

Cells were plated at 1000 cells per well into a 96-well plate with 5 technical replicates plated per condition. Cells were irradiated using a GammaCell 40 Irradiator (Best Theratronics). Non-irradiated plates were transported to the radiation facility, but not exposed. The 96-well plates were then immediately placed into the IncuCyte ZOOM live cell imaging system (Satroius). Phase-contrast images were taken every 4 hours for a total of 72 hours with 4 scan areas taken per well. Cell confluence was analyzed using the IncuCyte ZOOM analysis software. Percent confluence over time was used as a measure of longitudinal cell growth.

### Clonogenic Assay

Cells were plated at 500 cells per well into 6-well plates. The next day, cells were either left unirradiated (0 Gy) or irradiated with 4, 8, 12, or 16 Gy using a GammaCell 40 Irradiator (Best Theratronics). Media was changed 24 h after irradiation. Five days post-treatment, cells were washed in 1x PBS, before being fixed with 100% methanol for 10 minutes and stained with a 0.5% crystal violet solution for 30 minutes. Plates were imaged the next day on the LI-COR Odyssey near infrared imaging system. Clonogenic assay plate images were analyzed with ImageJ (Java 1.8.0_172), utilizing ImageJ’s thresholding, ROI (region of interest) manager, and analyze particles functions to count individual colonies.

### Immunofluorescence for **γ**H2AX

Cells were plated at 20,000 cells per well onto coverslips within the wells of 24 well plates. 24 hours post plating, cells were left un-irradiated (0 Gy) or irradiated with 3 Gy using a GammaCell 40 Irradiator (Best Theratronics). Coverslips were harvested and fixed with MeOH at 1, 6, 24, and 48 hours post-irradiation. Cells were then processed for immunofluorescence for γH2AX (1:500; Millipore, 05–636 JBW301). Secondary detection was accomplished using Alexa Fluor 488 goat anti-mouse IgG (1:500; Invitrogen, A28181). Nuclei were counterstained with Hoechst. Images were acquired using a Leica DM5500B upright epifluorescence microscope for a total number of 100–200 cells analyzed per n. Cells were scored as having more or less than 10 γH2AX foci.

### Western Blot analysis

For protein collection, plates were rinsed once with cold PBS, then cells were lysed with RIPA buffer plus cOmplete Protease Inhibitor Cocktail (Roche), PhosStop Phosphatase Inhibitor Cocktail (Roche), and PMSF. Cell lysates were left on ice for 10 minutes, vortexing every 2 minutes, then frozen at -80°C. Before use, lysates were thawed and spun at 16,000xg for 15 minutes at 4°C. Supernatant was collected and used for downstream analyses. Protein concentration was measured with the DC Protein Assay (Bio-Rad), following the manufacturer’s microplate assay protocol, and using an Infinite 200 PRO microplate reader (Tecan) to measure absorbance. Total protein lysate was combined with NuPAGE LDS Sample Buffer (Invitrogen) and NuPAGE Sample Reducing Agent (Invitrogen), then samples were heated for 10 minutes at 95°C. For *Nf1-/- DNp53* astrocyte and *Nf1-/- DNp53* Cas9 and p21 KD samples, 100 μg of protein was loaded, and samples were run on a NuPAGE 4-12% Bis-Tris Gel in MES SDS Running Buffer (Invitrogen). For wildtype mouse astrocyte and human GBM samples, 30 μg of protein was loaded, and samples were run on a NuPAGE 4-12% Bis-Tris Gel in MES SDS Running Buffer (Invitrogen). For FCG-Cas9 and FCG-Cas9 Nf1/p53 CRISPR samples, 75 μg of protein was loaded and samples were run on a NuPAGE 4-12% Bis-Tris Gel in MOPS SDS Running Buffer (Invitrogen). After separation by electrophoresis, proteins were transferred to Odyssey nitrocellulose membrane (LI-COR). Membranes were blocked for 1 hour at room temperature in Odyssey Blocking Buffer (LI-COR) or Intercept (PBS) Blocking Buffer (LI-COR) diluted 1:1 with PBS-T. Membranes were incubated overnight in primary antibody diluted in blocking buffer (1:1000 anti-cleaved caspase-3 Cell Signaling Technologies #9664, 1:4000 anti-α-Tubulin Sigma #T5168, 1:1000 anti-PARP Cell Signaling Technologies #9542, 1:2000 anti-mouse p21 abcam #ab188224, 1:1000 anti-Cdk2 Cell Signaling Technologies #2546, 1:1000 anti-human p21 Cell Signaling Technologies #2947, 1:200 anti-Nf1 Santa Cruz #sc-376886, 1:2000 anti-p53 Leica #NCL-L-p53-CM5p, 1:25,000 anti-Actin Sigma #A1978). Membranes were washed with PBS-T then incubated in secondary antibody diluted in blocking buffer for 1 hour at room temperature (1:30,000 IRDye 680RD Donkey anti-Rabbit or Donkey anti-mouse, 1:20,000 IRDye 800CW Donkey anti-Rabbit or Donkey anti-mouse (LI-COR)). Membranes were washed in PBS-T, then proteins were visualized using a ChemiDoc MP Imaging System (Bio-Rad). Band intensity was quantified using Image Lab Software Version 6.0.0 (Bio-Rad).

### Senescence-associated **β**-galactosidase staining

For *Nf1-/- DNp53* astrocytes and FCG GBM model astrocytes, cells were plated in 6 well plates at ∼70% confluence (70,000-100,000 cells per well depending on cell line/sex) and irradiated 24 hours later. 4 days after irradiation, cells were lifted, counted, and plated on poly-L-lysine (ScienCell) coated glass coverslips in 24 well plates at sub-confluency (15,000 cells per well). 24 hours later, cells were stained using the Senescence Associated β-galactosidase Staining Kit from Cell Signaling Technologies, according to the manufacturer’s instructions. Briefly, cells were washed once with PBS, then fixed for 15 minutes with provided Fixative Solution. After fixing, cells were rinsed twice with PBS, then incubated in β-galactosidase staining solution (pH 5.9-6.1) overnight (∼16 hours) in a 37°C dry incubator. The next morning, staining solution was removed and cells were rinsed twice with PBS. For mouse wildtype astrocytes, cells were grown in Primaria 6 well plates and irradiated at ∼70% confluence. 5 days after irradiation, cells were lifted, counted, and plated on poly-L-lysine coated glass coverslips in 24 well plates at sub-confluency (27,500 cells per well). 48 hours later, cells were stained using the Senescence Associated β-galactosidase Staining Kit. For human GBM lines, cells were grown in laminin coated 6 well plates and irradiated at ∼70% confluence. 4 days after irradiation, cells were lifted, counted, and plated on laminin coated glass coverslips in 24 well plates at sub-confluency (25,000 cells per well). 24 hours later, cells were stained using the Senescence Associated β-galactosidase Staining Kit. After SA-β-gal staining, cells were counterstained with Nuclear Fast Red (Vector Laboratories), following the manufacturer’s instructions. Coverslips were dehydrated then mounted on slides using Permount Mounting Medium (Fisher) diluted 2:1 with Xylene substitute. Images were taken on a Zeiss Axio Scope A1 upright light microscope at 10x magnification. Total cells and positive cells were counted by hand in ImageJ (FIJI Version 2.1.0) using the Cell Counter plugin. All images were taken and quantified by an experimenter blinded to the sex and experimental condition of the sample.

### Immunocytochemistry for Ki67

*Nf1-/- DNp53* astrocytes were plated in 6-well plates and irradiated the following day. 4 days after irradiation cells were trypsinized and counted, and 15,000 cells per well were plated on poly-L-lysine (ScienCell) coated glass coverslips in 24 well plates. 24 hours later cells were washed once with PBS, then fixed with 3.2% paraformaldehyde (Electron Microscopy Sciences) for 15 minutes. Cells were then washed twice with PBS and stored at 4°C until staining was performed. For Ki67 immunocytochemistry, cells were permeabilized with 0.2% Triton-X 100 for 15 minutes at room temperature and nonspecific antibody staining was blocked with antibody diluent (10% normal donkey serum, 1% bovine serum albumin (BSA) and 0.2% Triton-X 100 in PBS) for 1 hour at room temperature. Cells were then incubated in rabbit anti-Ki67 (Abcam ab15580, 1:500) or rabbit IgG control (Cell Signaling) overnight at 4°C, after which cells were washed and incubated in donkey anti-rabbit Alexa Fluor 555 (Invitrogen, 1:1000) for 1 hour at room temperature. Nuclei were stained with DAPI for 1 minute, then coverslips were mounted in Immu-Mount (Thermo Fisher Scientific). Images were taken on an Olympus BX60 (Olympus, Japan) fluorescence microscope at 10x magnification. Ki67 images were taken at 500 msec exposure. Color images were converted to 16bit and thresholded in ImageJ. Thresholds were determined using IgG controls. After thresholding, cell number was counted with the automated cell counter (Analyze Particles); anything <60 pixel^2^ was excluded from the count. The positive rate is the ratio of stained cells to total cells. All images were taken and quantified by an experimenter blinded to the sex and experimental condition of the sample.

### RNA isolation and cDNA preparation – Mouse GBM model astrocytes

RNA was isolated from *Nf1-/- DNp53* astrocytes and FCG GBM model astrocytes using TRIzol Reagent (Invitrogen), following the manufacturer’s protocol. Briefly, cells were grown in 10 cm dishes to ∼70% confluence, then irradiated. At desired timepoint after irradiation, media was aspirated, and cells were washed once with cold PBS. 1 ml of Trizol was added to the dish; cells were scraped into Trizol, allowed to sit 5 minutes at room temperature, then transferred to a 1.7 ml tube. 200 μl of chloroform was added, and samples were shaken vigorously for 15 seconds, then sat at room temperature for 3 minutes. Samples were spun at 12,000xg for 15 minutes at 4°C. The aqueous phase was transferred to a new tube, and 500 μl of isopropyl alcohol was added to precipitate RNA. Samples were incubated for 10 minutes at room temperature, then spun at 12,000xg for 10 minutes at 4°C. Supernatant was aspirated, and pellet was washed with 1 ml cold 75% ethanol. Samples were spun at 7500xg for 5 minutes at 4°C and ethanol was aspirated. After drying, pellet was resuspended in 35 μl molecular biology grade water (DNase/RNase free). RNA concentration was measured with a NanoDrop 1000 spectrophotometer (Thermo Scientific). RNA was treated with Amplification Grade DNAse I (Invitrogen) to eliminate genomic DNA, and cDNA was generated using the SuperScript III First-Strand Synthesis System (Invitrogen), according to the manufacturer’s instructions.

### RNA isolation and cDNA preparation – Mouse wildtype astrocytes

RNA was isolated from mouse wildtype astrocytes using the QIAGEN RNeasy Mini Kit, according to the manufacturer’s instructions. Briefly, astrocytes were grown in 10 cm Primaria dishes to ∼70% confluence, then irradiated. At desired timepoint after irradiation, media was aspirated, and cells were washed once with cold PBS. 350 μl of RLT buffer + β-mercaptoethanol was added and cells were scraped into RLT buffer, then transferred to a 1.7 ml tube. Cells were homogenized by passing through a 20-gauge needle attached to a 1 ml syringe ∼10 times, then 350 μl of 70% ethanol was added to the cell lysate and mixed by pipetting, before transferring to the RNeasy spin column. To eliminate genomic DNA, the optional on column DNase digestion was performed by adding 80 μl DNase I solution (QIAGEN) to the column membrane and incubating 15 minutes at room temperature. RNA was eluted by adding 30 μl molecular biology grade water (DNase/RNase free) directly to the column membrane, incubating at room temperature for 2.5 minutes, then spinning at 10,000xg for 1 minute. RNA concentration was measured with a NanoDrop 1000 spectrophotometer (Thermo Scientific). cDNA was generated using the QuantiTect Reverse Transcription Kit (QIAGEN) following the manufacturer’s instructions.

### Quantitative Real-time PCR

Quantitative RT-PCR was performed using gene specific primers and iTaq Universal SYBR Green Supermix (Bio-Rad). A standard curve was generated for each experiment by pooling cDNA from all samples and serially diluting to generate concentrations of 25 ng/μl, 12.5 ng/μl, 6.25 ng/μl, and 3.125 ng/μl. Samples were diluted to a concentration of 10 ng/μl. 2 μl of sample or standard was loaded in each well, and two technical replicates were run for each sample. A CFX Connect Real-Time PCR Detection System (Bio-Rad) was used to collect and analyze the amplification data. Analyses of standard curves, melt curves, and reverse-transcriptase-negative controls were used to verify primer and amplification reaction quality. Expression levels of genes of interest were normalized to the average expression of 3 housekeeping genes: *Gapdh*, *Actb, Rpl5.* Primer sequences: *Actb* (Forward: TGT ATT CCC CTC CAT CGT G, Reverse: CGC AGC TCA TTG TAG AA GG), *Ccnd1* (Forward: GTG CAT CTA CAC TGA CAA CTC, Reverse: TGG TCT GCT TGT TCT CAT CC), *Cdk2* (Forward: GCA CCA GGA CCT CAA GAA AT, Reverse: ACG GTG AGA ATG GCA GAA AG), *Cdkn1a* (Forward: AGA CAT TCA GAG CCA CAG GCA CCA, Reverse: GCA TCG CAA TCA CGG CGC AA), *Cdkn1b* (Forward: TGG TGG ACC AAA TGC CTG AC, Reverse: TTC GGG GAA CCG TCT GAA AC), *Cdkn2a* (Forward: GAC ATC GTG CGA TAT TTG CGT TCC G, Reverse: TTT AGC TCT GCT CTT GGG ATT GGC C), *Gapdh* (Forward: GGC AAA TTC AAC GGC ACA GT, Reverse: AGA TGG TGA TGG GCT TCC C), *Rpl5* (Forward: GGA AGC ACA TCA TGG GTC AGA, Reverse: TAC GCA TCT TCA TCT TCC TCC ATT), *Xist* (Forward: GCT GTA GTA GTC ACA GTC CCA, Reverse: CTG TGT TTG CCC CTT TGC TA).

### Statistics

Data were graphed and statistical analyses, including two-way ANOVAs, t-tests, one sample t-tests and dose response curves, were performed using GraphPad Prism software (Prism 9 for macOS Version 9.0.0). All tests were two-tailed unless otherwise specified. The two-way ANOVA models fit the effect of sex, time or dosage, and their interaction, followed by post-hoc multiple comparisons using the Bonferroni correction. Dose response curves were fit to normalized response versus dosage using the 4-parameter logistic regression dose-response model with the slopes estimated by maximizing the likelihood function. For *Nf1-/- DNp53* graphs normalized to the Male 0 Gy condition of the same cell line (Fig. 3b, d and Supplementary Fig. 2b, c, d), the Male 0 Gy values were derived by dividing by the Male 0 Gy average. Spearman rank-based correlation analysis between IC_50_ and gene expression was performed as described^7^ using R (version 3.5.0). Briefly, Spearman correlation coefficients of irradiation IC_50_ values with expression of either MC5 genes (17 genes), FC3 genes (9 genes), or randomly selected gene sets of the same size, were calculated. The Olkin-averaged Spearman correlation coefficients with IC_50_ across MC5 genes and across FC3 genes per cell line was summarized and tested against 0 with those derived for 1000 random gene sets serving as negative control. To explore the relationship between senescence and the various molecular markers focused on p16, p21, and p21/CDK2 ratio, linear regression analysis and Pearson’s correlation analysis was conducted using R (version 4.0.2). Pearson’s correlation coefficients (r) were calculated and the Fisher’s Z transformed coefficients were compared between sex by normal test. In the integrative linear regression modeling across sex, the dependent/outcome variable was senescence, as measured by SA-β-gal, while the independent/predictor variable includes the continuous mRNA or protein levels of the molecular marker of interest and sex. To assess the possible linear association between SA-β-gal and each of the continuous variables by sex, sex-specific intercepts and slopes were estimated from the integrative linear models. The slope difference between the male and female-specific slopes was derived with 95% confidence interval (CI) and tested against 0 by the 2-sided Wald test. Significance was claimed at p<0.05. Detailed slope and correlation estimation results can be found in Supplementary Table 3. For the *Nf1-/- DNp53* mRNA correlations only, mRNA expression levels from cells irradiated with 0, 6 or 9 Gy were correlated with SA-β-gal results from cells irradiated with 0, 6, or 8 Gy – thus resulting in a correlation consisting of 0, 6, and 8/9 Gy doses. For all other correlations, irradiation levels were exactly the same for protein/mRNA measures and SA-β-gal.

### Data Availability

The data that support the findings of this study are available from the corresponding author upon reasonable request.

## Acknowledgments

The authors would like to the thank the Department of Radiation Oncology, Washington University School of Medicine for use of the RS 2000 X-ray irradiator. This work was supported by NIH R01 CA174737-06 (JBR), NIH R21 NS098210 (JBR), Joshua’s Great Things (JBR), NIH R01 NS094670 (AHK), NIH R01 NS106612 (AHK), Siteman Investment Program Pre-R01 Research Development Award (AHK), NIH R03 CA227206 (MV), a Research Scholar Grant RSG-18-066-01-TBG from the American Cancer Society, funds from The Ohio State University Comprehensive Cancer Center/Department of Radiation Oncology (MV), and the National Institute of General Medical Sciences of the National Institutes of Health under award number 2T32GM068412-11A1 (MMT). LG and CMH were supported by the MARC U-STAR Program at Washington University in St. Louis, grant number T34 GM083914. JL also received support from the NCI Cancer Center Support Grant for the Siteman Cancer Center P30 CA091842 (PI: Dr. Timothy J. Eberlein).

