## Supplementary Materials for "Gonadal sex patterns p21-induced cellular senescence in mouse and human glioblastoma"

Supplementary Table 1. Molecular characteristics of patient tumors from which primary human GBM lines were derived

| Line | Sex | MGMT | Copy Number Gain | Copy Number Loss | Mutations | p53 Immuno-reactivity |
|---|---|---|---|---|---|---|
| B2 | Female | | *EGFR* | | *PTEN* | |
| B5 | Female | | *EGFR, BRAF, ALK* | | | |
| B18 | Female | Methylated | | | | |
| B30 | Male | Unmethylated | Polysomy 7 | | | |
| B31 | Male | Unmethylated | Polysomy 7 with *EGFR* amplification, *ALK* | | *ATM, APC* | |
| B36 | Male | Methylated | | 1p36 deletion/ monosomy 1, monosomy 10, 19q13 deletion | *NF1, PTEN, TERT* | Positive |
| B49 | Female | Unmethylated | *ALK* | Monosomy 10 | *PTEN, p53* | Positive |
| B51 | Female | | *EGFR* | 10q deletion/ monosomy 10 | | Positive |
| B66 | Male | Unmethylated | Polysomy 7 without *EGFR* amplification, Polysomy 8 | Monosomy 10, 19q deletion | *EGFR, PTEN, p53* | Positive |

Supplementary Table 2. MC5 and FC3 gene lists

| **MC5 Genes (17)** | **FC3 Genes (9)** |
|---|---|
| *BIRC5* | *AK5* |
| *CCNB1* | *AMIGO2* |
| *CCNB2* | *CHL1* |
| *CDC20* | *FERMT1* |
| *CKS2* | *IGFBP2* |
| *EZH2* | *PCDHB* |
| *KIF20A* | *PLAT* |
| *NEFH* | *POSTN* |
| *NEFM* | *SDC4* |
| *NES* | |
| *NUSAP1* | |
| *PBK* | |
| *PRC1* | |
| *PTTG1* | |
| *RRM2* | |
| *TOP2A* | |
| *TPX2* | |

Supplementary Table 3. Summary of correlation statistics

| Figure Panel | Cell Line | Sex | Variables | Slope Estimate (SE, p value) | Slope Difference F vs M (SE) | Slope Difference p value | Correlation (95% CI, p value) | Correlation Difference p value |
|---|---|---|---|---|---|---|---|---|
| 4a | Nf1-/- DNp53 | Male | SA-β-gal ~ *Cdkn2a* | 6.27 (6.68, 0.3562) | -9.60 (7.97) | 0.2395 | 0.36 (-0.19 to 0.73, 0.1931) | 0.1840 |
| | | Female | | -3.33 (4.36, 0.4520) | | | -0.17 (-0.63 to 0.38, 0.5479) | |
| 4b | Nf1-/- DNp53 | Male | SA-β-gal ~ *Cdkn1a* | 5.38 (2.40, 0.0335) | 1.99 (2.80) | 0.4823 | 0.59 (0.11 to 0.85, 0.0212) | 0.3464 |
| | | Female | | 7.38 (1.43, 0.00002) | | | 0.79 (0.46 to 0.93, 0.0005) | |
| 4c | Nf1-/- DNp53 | Male | SA-β-gal ~ *Cdkn1a/Cdk2* | 4.82 (2.37, 0.0524) | 3.38 (2.83) | 0.2428 | 0.53 (0.02 to 0.82, 0.0427) | 0.2011 |
| | | Female | | 8.20 (1.55, 0.00002) | | | 0.80 (0.50 to 0.93, 0.0003) | |
| 4e | Nf1-/- DNp53 | Male | SA-β-gal ~ p21/Cdk2 (24h) | 1.58 (1.87, 0.4072) | 3.48 (2.36) | 0.1526 | 0.27 (-0.28 to 0.69, 0.3356) | 0.2149 |
| | | Female | | 5.06 (1.43, 0.0016) | | | 0.65 (0.21 to 0.87, 0.0083) | |
| 4f | Nf1-/- DNp53 | Male | SA-β-gal ~ p21/Cdk2 (5d) | 2.59 (1.18, 0.0380) | 7.95 (2.38) | **0.0026** | 0.58 (0.09 to 0.84, 0.0244) | 0.3346 |
| | | Female | | 10.53 (2.07, 0.00003) | | | 0.78 (0.45 to 0.92, 0.0006) | |
| 5c | WT Astrocytes | Male | SA-β-gal ~ *Cdkn2a* | -32.44 (43.16, 0.4647) | 56.64 (45.66) | 0.2352 | -0.27 (-0.77 to 0.43, 0.4441) | 0.1418 |
| | | Female | | 24.20 (14.91, 0.1269) | | | 0.52 (-0.29 to 0.90, 0.1843) | |
| 5d | WT Astrocytes | Male | SA-β-gal ~ *Cdkn1a* | 5.52 (6.37, 0.4005) | 13.82 (9.40) | 0.1636 | 0.28 (-0.43 to 0.77, 0.4410) | 0.1839 |
| | | Female | | 19.34 (6.91, 0.0142) | | | 0.79 (0.18 to 0.96, 0.0207) | |
| 5e | WT Astrocytes | Male | SA-β-gal ~ *Cdkn1a/Cdk2* | 5.06 (3.43, 0.162) | 7.92 (4.73) | 0.1163 | 0.39 (-0.31 to 0.82, 0.2608) | **0.0243** |
| | | Female | | 12.98 (3.25, 0.0014) | | | 0.94 (0.70 to 0.99, 0.0005) | |
| 5g | WT Astrocytes | Male | SA-β-gal ~ p21 | 12.01 (9.25, 0.2034) | 18.19 (12.05) | 0.1409 | 0.33 (-0.30 to 0.76, 0.2909) | 0.2375 |
| | | Female | | 30.19 (7.73, 0.0005) | | | 0.67 (0.37 to 0.85, 0.0003) | |
| 5h | WT Astrocytes | Male | SA-β-gal ~ p21/Cdk2 | 1.33 (1.91, 0.4935) | 2.30 (2.32) | 0.3284 | 0.20 (-0.43 to 0.69, 0.5418) | 0.3304 |
| | | Female | | 3.63 (1.31, 0.0092) | | | 0.53 (0.16 to 0.77, 0.0081) | |
| 6c | Human GBM | Male | SA-β-gal ~ p21 (24h) | 6.57 (13.08, 0.6247) | 8.73 (13.53) | 0.5309 | 0.23 (-0.56 to 0.81, 0.5783) | 0.1116 |
| | | Female | | 15.30 (3.49, 0.0009) | | | 0.85 (0.35 to 0.97, 0.0080) | |
| 6d | Human GBM | Male | SA-β-gal ~ p21/Cdk2 (24h) | 3.77 (3.08, 0.2445) | -3.32 (3.09) | 0.3029 | 0.54 (-0.26 to 0.90, 0.1649) | 0.3159 |
| | | Female | | 0.45 (0.10, 0.0006) | | | 0.85 (0.35 to 0.97, 0.0081) | |
| 6e | Human GBM | Male | SA-β-gal ~ p21 (5d) | -5.50 (31.3, 0.8636) | 28.05 (31.79) | 0.3949 | -0.09 (-0.75 to 0.66, 0.8408) | **0.0461** |
| | | Female | | 22.56 (5.51, 0.0015) | | | 0.83 (0.29 to 0.97, 0.0115) | |
| 6f | Human GBM | Male | SA-β-gal ~ p21/Cdk2 (5d) | -1.69 (2.96, 0.5786) | 2.18 (2.96) | 0.4770 | -0.27 (-0.82 to 0.54, 0.5179) | **0.0192** |
| | | Female | | 0.48 (0.11, 0.0011) | | | 0.84 (0.32 to 0.97, 0.0099) | |
| 8d (left) | FCG GBM | XY+ | SA-β-gal ~ *Cdkn1a* | -3.81 (48.26, 0.9390) | N/A | N/A | -0.04 (-0.96 to 0.96, 0.9562) | N/A |
| | | XY- | | 86.62 (53.83, 0.1462) | | | 0.97 (0.11 to 1.00, 0.0315) | |
| | | XX+ | | 9.99 (41.46, 0.8157) | | | -0.12 (-0.95 to 0.97, 0.8797) | |
| | | XX- | | 73.31 (58.01, 0.2419) | | | 0.85 (-0.61 to 1.00, 0.1531) | |
| 8d (center) | FCG GBM | XY | SA-β-gal ~ *Cdkn1a* | 31.13 (33.22, 0.37) | -0.51 (42.73) | 0.9908 | 0.36 (-0.46 to 0.85, 0.386) | 0.9 |
| | | XX | | 30.63 (26.88, 0.28) | | | 0.42 (-0.40 to 0.87, 0.2968) | |

| | | | | | | | | |
|---|---|---|---|---|---|---|---|---|
| 8d (right) | FCG GBM | Sry+ | SA-β-gal ~ *Cdkn1a* | 10.66 (25.02, 0.6777) | 70.19 (41.51) | 0.1167 | 0.13 (-0.63 to 0.76, 0.7617) | **0.03** |
| | | Sry- | | 80.85 (33.13, 0.0311) | | | 0.91 (0.58 to 0.98, 0.0016) | |
| 8e (left) | FCG GBM | XY+ | SA-β-gal ~ *Cdkn1a/Cdk2* | -19.29 (39.83, 0.6412) | N/A | N/A | -0.26 (-0.98 to 0.93, 0.7402) | N/A |
| | | XY- | | 121.83 (75.69, 0.1462) | | | 0.94 (-0.24 to 1.00, 0.0625) | |
| | | XX+ | | -24.23 (35.11, 0.5096) | | | -0.33 (-0.98 to 0.92, 0.6665) | |
| | | XX- | | 106.28 (79.69, 0.2190) | | | 0.87 (-0.57 to 1.00, 0.1348) | |
| 8e (center) | FCG GBM | XY | SA-β-gal ~ *Cdkn1a/Cdk2* | 7.21 (36.51, 0.8468) | -1.56 (46.98) | 0.974 | 0.08 (-0.66 to 0.74, 0.848) | 0.9941 |
| | | XX | | 5.65 (29.58, 0.8518) | | | 0.08 (-0.66 to 0.74, 0.8567) | |
| 8e (right) | FCG GBM | Sry+ | SA-β-gal ~ *Cdkn1a/Cdk2* | -16.42 (22.02, 0.4702) | 131.06 (51.73) | **0.0262** | -0.22 (-0.80 to 0.57, 0.5964) | **0.01** |
| | | Sry- | | 114.64 (46.81, 0.0306) | | | 0.91 (0.55 to 0.98, 0.002) | |

*N/A = Not applicable

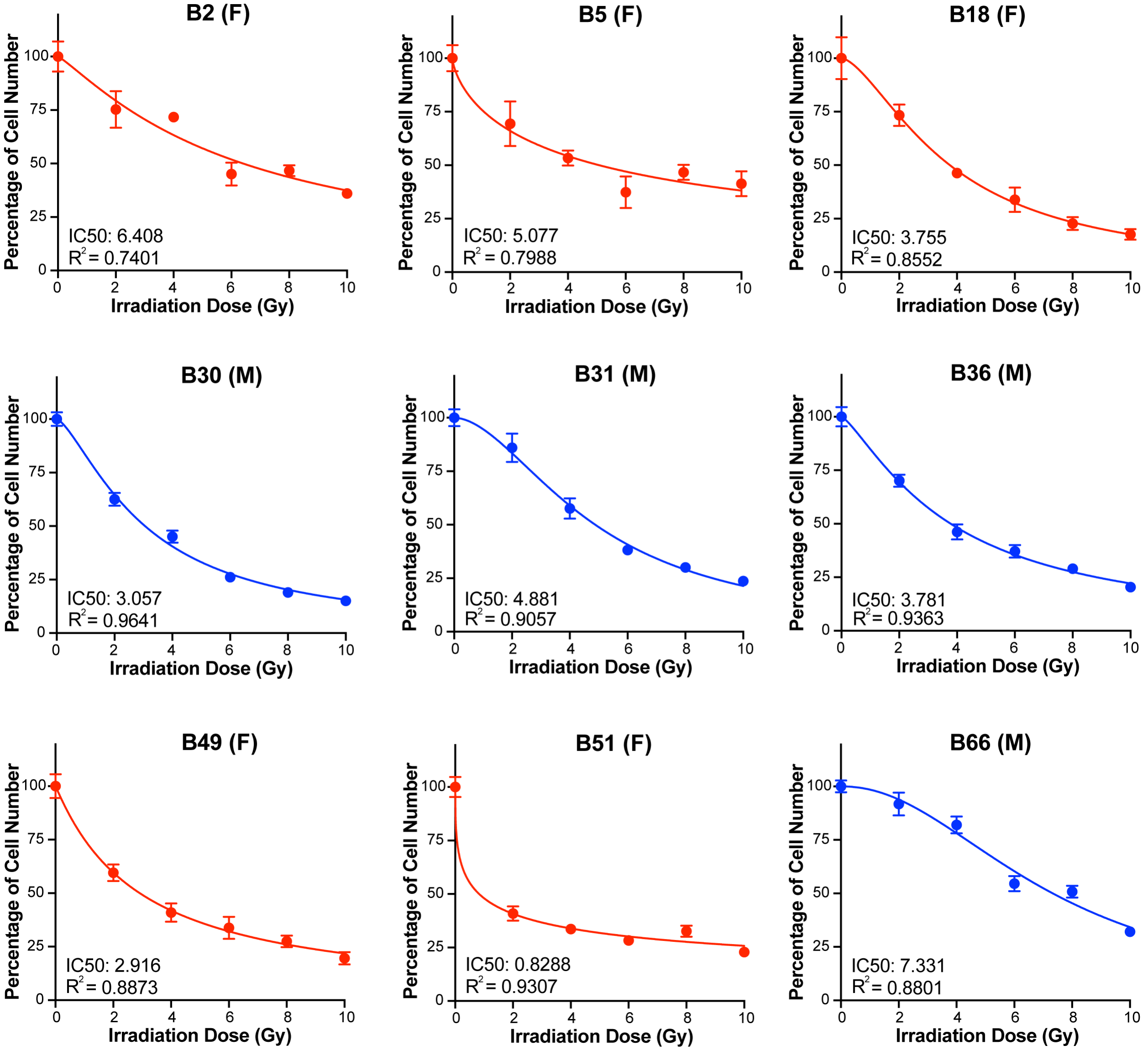


**Supplementary Fig. 1: Irradiation dose response curves for male and female human GBM lines.** Irradiation dose response curves for 4 male (M) and 5 female (F) primary human GBM lines. Cells were irradiated 24 hours after plating, and cell number was counted on Day 4 after irradiation. Percentage of cell number was calculated by dividing the cell counts for each dose by the average cell number at 0 Gy. Data are means +/- SEM (B5 n=3/dose, All other lines n=6/dose).

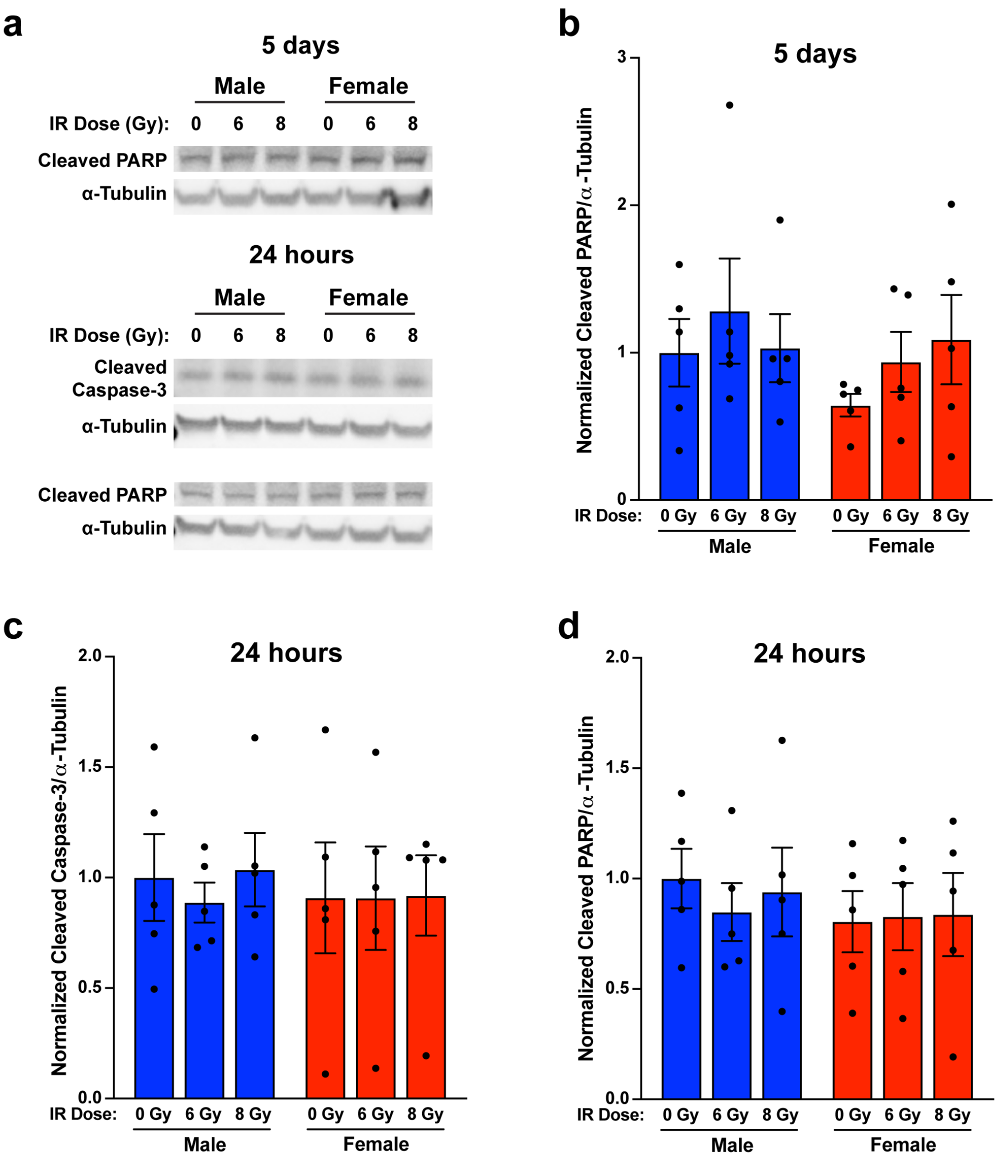


**Supplementary Fig. 2: Apoptosis is not the primary response to irradiation at either early or late timepoints. a** Representative western blot images of cleaved PARP at 5 days after irradiation, and cleaved caspase-3 and cleaved PARP at 24 hours after irradiation. Representative bands from one male and one female *Nf1-/- DNp53* cell line are shown. **b** Quantification of cleaved PARP levels measured by western blot 5 days after irradiation. Values were normalized to the corresponding Male 0 Gy condition of the same cell line, which was arbitrarily set at 1. Two-way ANOVA: Dose p=0.4664, Sex p=0.3622, Interaction p=0.7214. Data are means +/- SEM (n=5/sex/dose). **c** Quantification of cleaved caspase-3 levels measured by western blot 24 hours after irradiation. Values were normalized to the corresponding Male 0 Gy condition of the same cell line, which was arbitrarily set at 1. Two-way ANOVA: Dose p=0.8523, Sex p=0.7224, Interaction p=0.9131. Data are means +/- SEM (n=5/sex/dose). **d** Quantification of cleaved PARP levels measured by western blot 24 hours after irradiation. Values were normalized to the corresponding Male 0 Gy condition of the same cell line, which was arbitrarily set at 1. Two-way ANOVA: Dose p=0.9142, Sex p=0.4255, Interaction p=0.8619. Data are means +/- SEM (n=5/sex/dose).

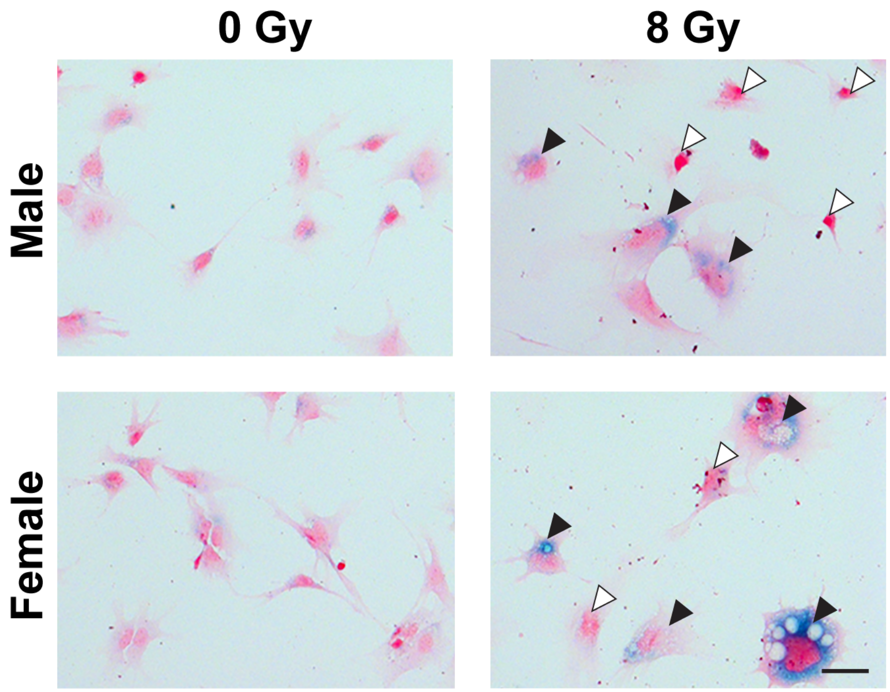


**Supplementary Fig. 3: Irradiation induces changes in cell size and shape consistent with senescence.** Example images of male and female *Nf1-/- DNp53* astrocytes stained for SA-β-gal (blue) 5 days after irradiation with 0 or 8 Gy, followed by counterstaining with nuclear fast red (pink). Scale bar, 50 μm. Images are magnified from Fig. 3c. Untreated male and female cells show some variation in size and morphology but are generally small with very little blue staining. Treated cells show much greater variation in size and morphology, and an increase in the percentage of cells with blue staining. Black arrowheads indicate large, irregularly shaped cells that are present after irradiation; white arrowheads indicate cells that retain normal *Nf1-/- DNp53* astrocyte morphology following irradiation.

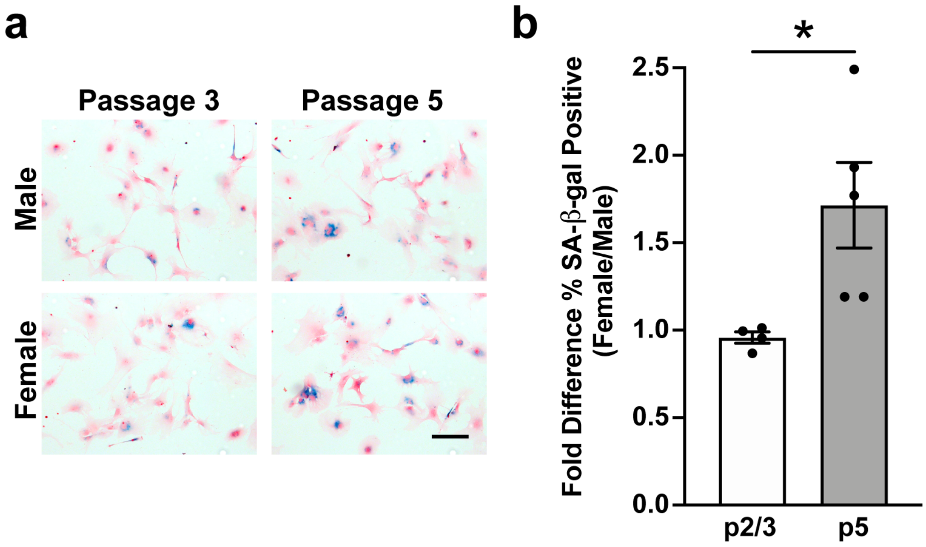


**Supplementary Fig. 4: Female wildtype astrocytes have increased senescent cells at later passages. a** Example images of male and female wildtype mouse astrocytes at low passage (p3) or high passage (p5) stained for SA-β-gal and counterstained with nuclear fast red. Scale bar, 150 μm. **b** Fold difference in the percentage of SA-β-gal positive cells in male and female cultures at either low (p2/p3) or high (p5) passages. *$p<0.05$, two-tailed t-test with Welch's correction. Data are means +/- SEM (n=4-5/group).

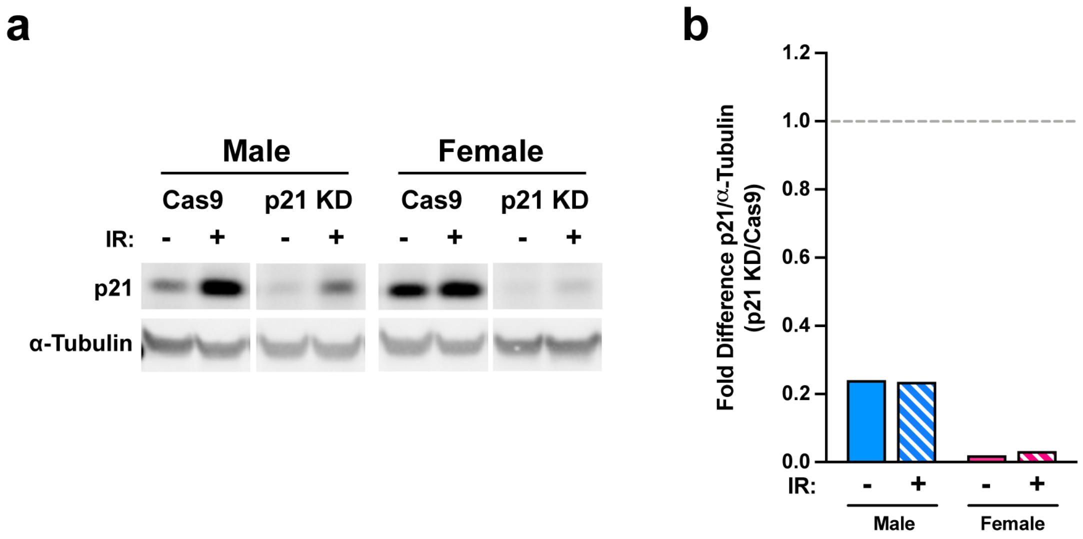


**Supplementary Fig. 5: Confirmation of p21 knockdown. a** Western blot images of p21 levels in *Nf1-/- DNp53* Cas9 control and p21 knockdown male and female astrocytes 24 hours after irradiation with 0 or 8 Gy. **b** Fold knockdown in p21 was calculated by dividing the levels of p21 expression in p21 KD lines by the levels in Cas9 control lines under non-irradiated and irradiated conditions.

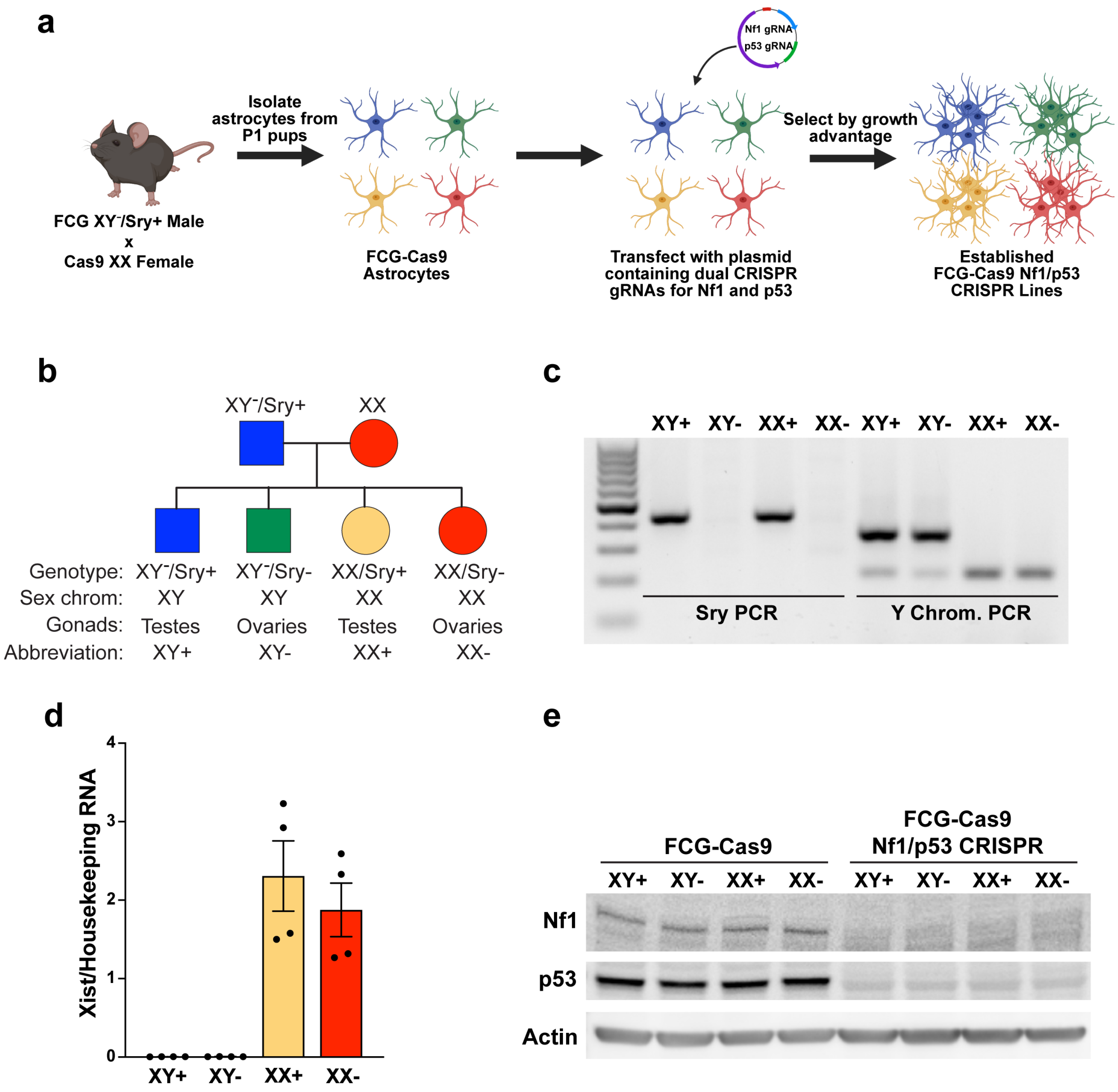


**Supplementary Fig. 6: Four Core Genotypes model of GBM.** **a** Schematic for generation of the FCG GBM model (created with BioRender.com). **b** Diagram of the four genotypes resulting from the FCG mouse model. **c** Genotyping PCR confirming correct genotypes in astrocyte cultures isolated from FCG-Cas9 postnatal day 1 pups. **d** Expression levels of the lncRNA Xist, measured by qPCR, in FCG-Cas9 Nf1/p53 CRISPR astrocytes, showing appropriate expression in XX cells only, regardless of gonadal sex. Results are from 2 cell lines, with 2 technical replicates each. Data are means +/- SEM (n=4/genotype). **e** Western blot images of Nf1 and p53 expression in FCG-Cas9 and FCG-Cas9 Nf1/p53 CRISPR astrocytes, confirming knockdown of these two proteins in the FCG GBM model.
